# Engineering a three-dimensional multilayer multicellular model of endometrial cancer for high throughput drug screening and novel treatment methods

**DOI:** 10.1101/2024.02.20.581239

**Authors:** Ines A Cadena, Claire Rowlands, Mina R Buchanan, Molly A Jenne, Bailey Keefe, Alyssa Almer, Ndubuisi Obasi, Conor G Harris, Willie E Rochefort, Brittany E. Givens, Kaitlin C Fogg

## Abstract

Endometrial cancer is one of the most common gynecological cancers in the world, with an estimated 382,000 new cases and 90,000 deaths each year. There is no specific treatment, as the underlying causes of endometrial cancer neoplasia are poorly understood. This study focuses on the development and validation of a three-dimensional (3D) *in vitro* multilayer, multicellularhydrogel that facilitates drug screening analysis. We hypothesized that a specific combination of natural (collagen type I and IV, fibrinogen, fibronectin, Laminin) and synthetic (GELMA, PEGDA) polymers would maximize microvessel formation and cell invasion. The 3D model incorporates human microvascular endothelial cells (hMVEC) and endometrial cancer cells (HEC-1A) atop hydrogel formulations mimicking cell-specific extracellular matrix components. Using a D-optimal experimental design, 45 hydrogel combinations were generated. The predicted hydrogel formulation to maximize all cell responses enhanced higher microvessel formation and cancer invasion compared to the gold standard Matrigel. Subsequent validation emphasizes the importance of a disease-specific model and cell crosstalk in maximizing microvessel formation and cancer invasion. The optimized 3D model adeptly captures variances in cell responses among endometrial cancer cell lines from distinct stages. Finally, the platform is employed to compare cell viability, microvessel formation, and cancer invasion across Ishikawa, KLE, and HEC-1A cells after Paclitaxel exposure, delivered both as a free drug and loaded in poly(caprolactone) (PCL) nanoparticles. Overall, this study provides a valuable tool for exploring intricate interactions within the tumor microenvironment, offering a holistic understanding of cell responses and fostering the development of targeted therapeutic strategies for endometrial cancer.

## 1. Introduction

Endometrial cancer, the most common gynecological cancer in developed countries and the sixth most common type of cancer worldwide, presents a significant healthcare challenge.^1,2^ In 2023, 66,200 women were diagnosed with this disease and over 13,000 women died from endometrial cancer in the United States alone.^3^ While early-stage endometrial cancer can be effectively treated with surgery, chemotherapy, and radiation, advanced stages often have poor prognoses, with a five year survival rate under 20%, primarily due to metastasis.^4^ Despite increasing incidence rates in the United States, treatment advancements have been limited, with no significant FDA-approved therapies since the introduction of progestin therapies in the 1970s followed by approval of pembrolizumab and lenvatinib for recurrent or metastatic endometrial cancer in 2019.^5^ Notably, the combination of carboplatin and Paclitaxel is a standard front-line treatment for advanced endometrial cancer, yet it is not specifically listed as an FDA-approved drug for endometrial cancer.^6^ Furthermore, response rates to Paclitaxel remain low, with average reponse rates of 62% and overall survival times of less than three years.^7^ Thus there is an urgent need for new therapeutic options for endometrial cancer patients.

An improvement to paclitaxel therapeutics is the nanoparticle formulation. An albumin-bound paclitaxel nanoaparticle (nab-paclitaxel, Abraxane®) shows enhanced efficacy in patients who have previously experienced resistance to other chemotherapeutics. Similar to paclitaxel, nab-paclitaxel is not specifically FDA-approved for treating endometrial cancer. Nanoparticle formulations show enhanced clinical efficacy due to the ehanced permeability and retention (EPR) effect. EPR occurs in solid tumors and is caused by leaky vasculature allowing particles to easily enter the tumor environment, but due to ineffective lymphatic drainage, the particles have increased retention.^8^ In addition to the EPR effect, naparticle formulations mitigate limitations in bioavailbility, solubility, and absorption compared with free drug.^9,10^ A common way to create these nanoformulations is to encapsulate the drug in a polymeric nanoparticle. These formulations can be commonly referred to as drug delivery systems (DDSs). These DDSs often have a sustained release profiles which can increase the efficacy of the formulation by maintaining a consistent drug concentration in the tumor environment.^11^ Poly (caprolactone) (PCL) is an aliphatic polyester that is biodegradable, biocompatible, and has been FDA approved for use in the human body ranging from implanted devices to sutures. It is also a bioresorbable polymer as it can be completely degraded *in vivo* primarily by hydrolysis.^12^ Although not as common for DDS as some of its other aliphatic polyester counterparts such as poly(lactic-co-glycolic acid) (PLGA) or poly(lactic acid) (PLA), it remains a promising polymer for drug delivery due to being highly tunable with diffusion dominating short-term drug release as well as low production cost.^9^ Creating Paclitaxel loaded PCL nanoparticles will reduce the current limitations of free Paclitaxel such as its low solubility which will create a more effective treatment option.^13,14^

As novel treatment strategies are developed, the need for physiologically relevant and high-throughput screening platforms has become increasingly abundant. High-throughput screening is widely utilized for quickly identifying compounds with therapeutic potential. However, the quality of this screening depends upon the relevance of the model to the desired therapeutic target. Recent advancements in 3D *in vitro* models for endometrial cancer have led to more accurate representation of the tumor microenvironment compared to traditional 2D models or Matrigel. ^15,16^ Notably, a novel hydrogel derived from decellularized endometrium demonstrated that incorporating ECM components native to the endometrium increased reproducibility of endometrial organoids compared to Matrigel.^17^ Additionally, an innovative disintegration-controllable supramolecular gelatin hydrogel was used to develop a co-culture system of endometrial tumor spheroids and tumor-associated macrophages, shedding light on the interplay between M2 macrophages, ECM components, and endometrial cancer cells.^18^ These studies underscore the importance of including endometrial cancer-specific ECM components in 3D models and using 3D models for high-throughput screening. However, microvessel formation is a critical step in endometrial cancer progression and these models did not include endothelial cells, which is a clear area for improvement addressed in our model. Additionally, the integration of these models with high-throughput screening remains a challenge, limiting their application to drug screening and large-scale mechanistic studies. Thus, an overarching goal of this paper was to develop a novel platform that captures 3D metastatic behavior and interfaces with standard high throughput drug screening methods.

Building on our previous cervical cancer model^19^ and using a Design of Experiment (DOE) approach, we bridged the gap between classic tissue engineering models of cancer and high throughput screening methods for drug development. We created a multilayer multicellular platform that incorporated ECM components reflective of the tumor microenvironment. This model not only supported endometrial cancer cell lines from different endometrial cancer stages but also demonstrated enhanced performance comparted to Matrigel, the current gold standard for 3D drug screening.^20,21^ We then used this 3D platform to perform a comprehensive analysis comparing dose-response curves in 2D and 3D models treated with Paclitaxel and examining the effects of Paclitaxel delivered as a free drug and in PCL nanoparticles. The PCL nanoparticles effectively reduced cell viability compared with the free drug, showcasing the model’s potential in evaluating novel drug delivery systems. This study highlights the power of our 3D in vitro model as a versatile tool for in-depth exploration within the endometrial tumor microenvironment, offering critical insights for targeted therapeutic development. Furthermore, we are evaluating free Paclitaxel as a well-understood drug treatment in endometrial cancer and a novel nanoparticle-encapsulated form of Paclitaxel.

## 2. Materials and Methods

### Cell lines and reagents

Unless stated, all reagents were purchased from ThermoFisher (Waltham, MA). Human microvascular endothelial cells (hMVEC) were purchased from Lonza (hMVEC 33226, Walkersville, MD) and used without additional characterization. Cells were expanded in EGM-2 MV media (EBM-2 supplemented with Lonza’s SingleQuot supplements: hydrocortisone, human basic fibroblast growth factor (FGF2), human vascular endothelial growth factor (VEGF), human insulin-like growth factor (IGF), human epidermal growth factor (EGF), ascorbic acid, and gentamycin) and further supplemented with 5% fetal bovine serum (FBS) until use at passage 5. Human endometrial cancer cell lines HEC-1A (ATCC HTB-112™) and KLE (ATCC CRL-1622) and the human cervical cancer cell line SiHa (ATCC® HTB-35™) were purchased from ATCC (Manassas, VA) and used without additional characterization. The cells were cultured in McCoy’s 5A medium (HEC-1A, ATCC), DMEM: F12 medium (KLE, ATCC), and Eagle’s Minimum Essential Medium (SiHa, EMEM, ATCC), supplemented with 1%penicillin–streptomycin (Sigma-Aldrich, St. Louis, MO, USA) and 10% FBS. Ishikawa H cells were gifted from Dr. Aliasger Salem (University of Iowa, Iowa City, IA) to Dr. Brittany Givens. Ishikawa cells were cultured in DMEM medium (Quality Biological, Gaithersburg, MD) supplemented with 10% FBS and 1%penicillin– streptomycin. HEC-1A and KLE cells are categorized as Type II endometrial cancer cell lines, known for their highly aggressive and invasive cancer characteristics. In contrast, Ishikawa cells are identified as a Type I endometrial cancer cell line, characterized by low histological differentiation, representing an early-stage disease.[4] Furthermore, both the Hec-1A and the Ishikawa cell lines are considered Paclitaxel-insensitive, whereas KLE cells are considered Paclitaxel-sensitive.^22–25^ All cell types were expanded in standard cell culture conditions (37°C, 21% O_2_, 5% CO_2_) and subcultured before they reached 80% confluency.

### Evaluation of phenotypic cell response

All reagents were purchased from ThermoFisher (Waltham, MA) unless stated. Cell phenotypic response was monitored by staining hMVEC cells, endometrial cancer cells, and cervical cancer cells with fluorescent dyes before the experiment. CellTracker Green CMFDA (C7025), CellTracker Blue CMF2HC (C12881), CellTracker Red CMTPX (C34552) or CellTracker Deep Red (C34565) were used to visualize cells over time. From these images, we calculated endothelial cell microvessel formation, cancer cell invasion, and cell coverage over time. For all experiments, two-channel 100 µm z-stack images were taken every 3 hours for 48 hours using a Cytation 5 cell imaging multimode reader (Agilent Technologies). Images were processed with NIH Fiji-ImageJ and cell coverage was measured for each cell type at every time point by calculating the area within the well covered by cells and dividing it by the total well area. Cell invasion depth was measured using Gen 5 software (Agilent Technologies), and endothelial microvessel formation was quantified by measuring the average microvessel length at each time point with Fiji (NIH, Bethesda, MD).

Confocal microscopy images were taken for endometrial cancer cell lines seeded on 0.01% poly(L-lysine)-coated cover slips (Sigma Aldrich, St. Louis, MO). Cells were incubated for 24 hours prior to fixation. Cells were fixed with 4% paraformaldehyde for 10 minutes followed by 0.1% Triton-X-100 (Research Products International Corp) in 1X DPBS for five minutes. Non-specific binding was blocked with 10 mg/mL bovine serum albumin (BSA, Sigma Aldrich) in 1X DPBS for 20 minutes. Cells were stained with 0.165 uM AlexaFluor488-phalloidin (ThermoFisher) for 20 minutes, and then mounted onto coverslips using ProLong Diamon Anitfade Mountant with DAPI (ThermoFisher) sealed with clear nail polish. Cells were imaged on an Upright Zeiss LSM 880 multi-photon microscope.

Cell viability was quantified models using cell titer-glo 3D cell viability assay (Promega, Madison, WI). Following the manufacturer’s instructions, the reagent was added on top of the wells at either 24 or 48 hours post dosage of the chemotherapy agent. Cell metabolic activity was quantified in 2D models using an MTT assay (Thermo Fisher, Waltham, MA). For the MTT assays, all three endometrial cancer cell lines were seeded at 1.5 x 10^4^ cells/well in a 96-well plate. The cells were allowed to incubate overnight before adding the chemotherapy agent as free Paclitaxel or PCL particles loaded with Paclitaxel. At 24 or 48 hours, the media was removed and replaced with 100 µL of media as well as 10 µL of 5 mg/mL MTT in 1X Dulbecco’s Phosphate Buffered Saline (DPBS). The cells were then incubated for four more hours. Finally, the media was removed and 100 µL of an isopropanol:dimethylsulfoxide (IPA:DMSO, 9:1) solution was added. Absorbance was then measured at 570 nm on BioTek Synergy H1 plate reader.

### Multilayer Hydrogel Fabrication

We have previously developed a multilayer multicellular 3D *in vitro* model of cervical cancer that can support cervical cancer invasion and endothelial cell microvessel formation over 48 hours of culture.^26^ Using a similar approach, we developed a novel multilayer multicellular model of endometrial cancer (**Fig. 1A**). Constructs were fabricated in specialized µ-Plate Angiogenesis 96-wells (ibidi, Munich, Germany) by layering 10 µL of the bottom hydrogel formulation into each well, and 9,400 CellTracker Green labeled hMVEC cells in 40 µL of EGM-2 MV were pipetted on top of each hydrogel. The endothelial cells were allowed to attach for four hours at 37°C and 5% CO_2_ (**Fig. 1B, 1C**). The media was then removed, and 25 µL of the top hydrogel formulation was pipetted on top of the endothelial cells. Finally, CellTracker Red labeled endometrial cancer cells were seeded on top of each gel at 12,500 cells in 25 µL of media, with a 1:1 ratio of EGM-2 MV and cancer cell medium (**Fig. 1D, 1E**). The media was changed every 12 hours, and cells were imaged every 3 hours for 48 hours using a Cytation 5 cell imaging multimode reader (Agilent Technologies) .

**Figure 1.**
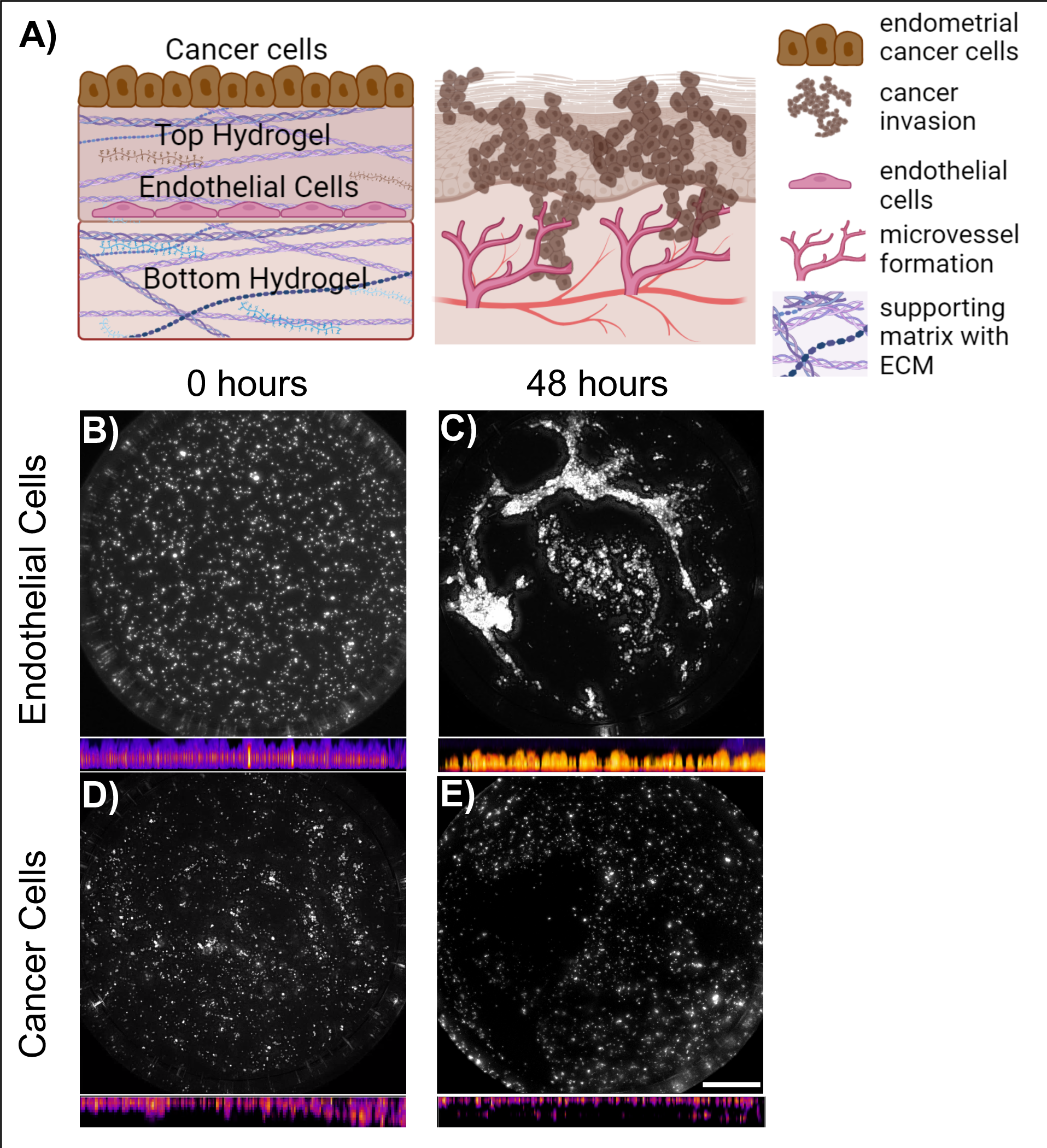
Multilayer multicellular model for endometrial cancer. (**A**) Schematic diagram of 3D endometrial cancer model (created with BioRender.com). 2D and 3D visualization of human microvascular endothelial cells (hMVEC) and endometrial cancer cells (HEC-1A). At time 0 hours (**B and D**) and 48 hours after seeded (**C and E**). 3D visualization created using FIJI (NIH). Scale bar 1000 µm

To prepare the bottom hydrogel, stock solutions of bovine collagen type I (10 mg/ml, FibriCol, Advanced BioMatrix), fibrinogen from human plasma (30 mg/mL Sigma-Aldrich), 50 U/mL thrombin (20 U, Sigma-Aldrich) and either polyethylene glycol diacrylate (PEGDA) (11.7 %w/v, Advanced BioMatrix, Carlsbad, CA, USA) or gelatin methacryloyl (GelMA) (8.7% w/v, Advanced BioMatrix) were mixed and photo-crosslinked using a CL-1000 UVP crosslinker with irgacure 2959 (365nm) as specified by manufactured (10% w/v, Advanced BioMatrix). 10µL of the final gel formulation was added to each well and the plate was incubated for one hour at 37°C. The top layer hydrogel was fabricated by combining the stock concentrations of laminin (1 mg/mL, Sigma-Aldrich),**^B^**h**^)^**uman fibronectin (1 mg/mL, **^C^**Ad**^)^**vanced Biomatrix), collagen IV, human (1 mg/mL, Advanced Biomatrix) and either GelMA (8.7% v/w) or PEGDA (11.7% w/v). After carefully removing the media, 25 uL of the final hydrogel formulation was added on top of the hMVEC layer. The construct was photo-crosslinked for 30 seconds in a CL-1000 UVP crosslinker using the same conditions as above. The plate was incubated for one hour at 37°C. Then, labeled endometrial cancer cells were added to the hydrogels and covered with 10 µL of media, with a 1:1 ratio of EGM-2 MV and cancer cell medium. The pH was adjusted for all the hydrogel formulations by adding sodium hydroxide (NaOH, 0.1N, Sigma-Aldrich), 1x PBS and 10x PBS (Research Products International Corp, Mount Prospect, IL, USA). As a control, multilayer Matrigel constructs were made using the same methods described above except replacing the custom hydrogel formulations with Growth Factor Reduced Basement Membrane Matrigel in both layers (9.2 mg/mL protein concentration, Corning, MA, USA).

### Design of Experiments

We determined the significance and interactions of the concentrations of natural and synthetic polymers present in each hydrogel layer on the function of each cell type. A D-Optimal design with quantitative and qualitative factors was created with MODDE Pro software version 12.1 (Sartorius AG, Göttingen, Germany) (**Supplemental Table 1**). Five quantitative factors were selected. The concentration of collagen type I was varied from 0.5 to 2.5 mg/mL, fibrinogen from 0.5 to 2.5 mg/mL, collagen type IV from 0.01 to 0.2 mg/mL, fibronectin from 0.125 to 0.175 mg/mL, and laminin from 0.5 to 2 µg/mL, all in 1X PBS. The two qualitative factors were the choice of either PEGDA (10% w/v) or GelMA (7%w/v) in both top and bottom hydrogel layers. The output variables were endothelial cell microvessel length, endothelial cell coverage, endometrial cancer invasion depth and endometrial cancer coverage. We used the HEC-1A endometrial cancer cell line to develop and validate the models as they could be used for further drug sensitivity studies. The D-Optimal design generated 45 unique hydrogel combinations with a centrally repeated condition to optimize the Design’s G-efficiency (**Supplemental Table 2**), which assessed how well the D-Optimal design compared to a fractional factorial design. The significance and interaction of all the input variables on the measured output variables were evaluated based on the R^2^ and Q^2^ diagnostics of the fit. The model was validated by evaluating the hydrogel formulations that were predicted to either maximize or minimize all of the phenotypic cell responses simultaneously (**Supplemental Table 3**).

To further explore the importance of having a multilayer system on cell interactions and responses, we compared the optimized multilayer model predicted by DOE to maximize all cell responses in monoculture and co-culture models. We seeded hMVEC cells on top of the optimized hydrogel formulation and the endometrial cancer cell line HEC-1A on top of the optimized top hydrogel formulation. We measured phenotypic cell responses for 48 hours. Differences between the monoculture and co-culture models were assessed by measuring changes in microvessel formation, invasion depth, and cell area over time. Matrigel was used as a control, replacing both of the hydrogel formulations.

### Comparing endometrial and cervical cancer-specific three-dimensional models

To explore the importance of building a 3D *in vitro* model specific to endometrial cancer, we compared our optimized multilayer model for endometrial cancer with our cervical cancer multilayer model that we have previously published.^26^ We measured differences in phenotypic cell responses with cervical cancer cells (SiHa, CellTracker Deep red labeled) and endometrial cancer cells (HEC-1A, CellTracker Deep red labeled) co-cultured on top of human microvascular endothelial cells (hMVEC, Cell Tracker green labeled) cells. Specifically, we compared changes in microvessel formation and invasion when cancer cells were seeded on their specific matrix and when they were seeded on the opposite models. A schematic of the experimental design is shown in **Figure 2**. Matrigel was used as a control, replacing the hydrogel formulations at the top and bottom.

**Figure 2.**
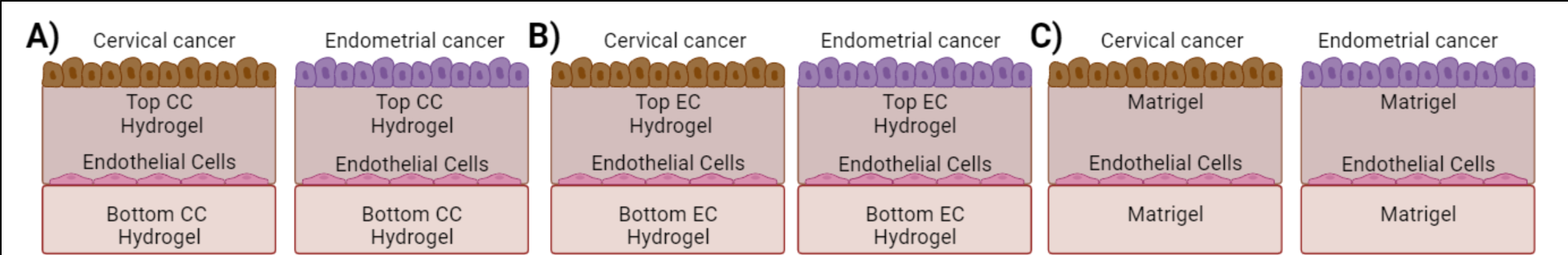
Schematic of experimental design for cervical cancer and endometrial cancer models. Multilayer multicellular models of cervical cancer (C.C.) and endometrial cancer (E.C.) were prepared with human microvascular endothelial cells (hMVEC) and either cervical cancer cells (SiHa) or endometrial cancer cells (HEC-1A). Phenotypic responses were compared when cells were seeded on optimized hydrogel formulations for (**A**) cervical cancer (C.C.), (**B**) endometrial cancer (E.C.), or (**C**) Matrigel.

### Assessment of material properties

Dynamic oscillatory shear measurements were used to evaluate the rheological properties of the bottom and top hydrogels of the optimized hydrogel formulation, as well as GFD Matrigel. The hydrogels with GelMA or PEGDA were photo crosslinked with 365 nm U.V. light and incubated at 37°C before rheological testing. Using the previously demonstrated method,[3] we use an AR-G2 rheometer (T.A. Instruments, New Castle, DE) equipped with a 20 mm standard steel parallel top plate and a bottom standard Peltier plate to conduct a rheological characterization of the hydrogels. Due to the tendency for the gels to slip, 150 grit (120 µm particle size) sandpaper was adhered to both the top and bottom fixtures. The gap was adjusted between measurements to ensure contact between the top plate and the gel such that the normal force was 0.1 N. All measurements were performed at a constant temperature of 37 °C using a raised Peltier plate control system. Frequency sweeps were conducted from 0.1 to 10 rad/sec within the linear viscoelastic (LVE) region at a constant strain amplitude of 5%. The storage moduli (*G*’) and loss moduli (*G*“) were measured across the frequency range, the average of the linear region was calculated for each hydrogel, and three consecutive technical replicates were conducted for each hydrogel formulation. All the hydrogel formulations were kept at 37°C in the incubator before the reading. The gels were allowed to equilibrate to the plate temperature of 37°C for 15 minutes, at which point frequency sweeps were conducted.

### Paclitaxel-loaded nanoparticles preparation

Paclitaxel-loaded poly(caprolactone) (PCL) nanoparticles were prepared using an emulsion solvent evaporation (o/w) method. The oil phase consisted of 60 mg of PCL (Mn 80,000, Sigma Aldrich, Milwaukee, WI), 3 mg of Paclitaxel (Selleck Chemicals LLC, Houston, TX), and 3 mL of dichloromethane (DCM) (VWR, Radnor, PA). The water phase consisted of a 2.5% (w/v) of poly(vinyl alcohol) (PVA) (87-90% hydrolyzed, average molecular weight 30,000-70,000, Sigma-Aldrich, Milwaukee, WI) in deionized water. The water phase was stirred for 20 minutes to allow the PVA to dissolve. The oil phase was combined and sat for 15 minutes until PCL and Paclitaxel were dissolved and vortexed prior to sonication. For the sonication step, 50 mL of the water phase was used. While the probe sonicator (Q500 Sonicator, QSonica, Newton, CT) was running, the oil phase was slowly added dropwise directly onto the probe. The sonicator ran at 50% power for 75s. After sonication, the emulsion was added to 30 mL of the water phase and stirred at 350 rpm for 30 minutes to allow the DCM to evaporate. The solution was then centrifuged at 1000 rcf at 20°C for 10 minutes. The supernatant was collected and spun again at 3000 rcf for 10 minutes. The pellet formed at 1000 rcf waws discarded. The pellet formed at 3000 rcf was washed three times with DI water at 3000 rcf to remove the PVA. After washing, the particles were suspended in DI water and frozen at -20°C then lyophilized (FreezeZone Bench Top freeze dryer, Labconco, Kansas City, MO) for at least 24 hours or until only a fine powder remained. The particles were stored at -20°C until use.

### Dose-response analysis of Paclitaxel

To compare free-drug and nanoparticle-loaded Paclitaxel, we conducted a dose-response analysis of Paclitaxel in two-dimensional (2D) cultures (endometrial cancer cells seeded on tissue culture plates) and three-dimensional models (using our co-culture 3D *in vitro* hydrogel model). Free-drug Paclitaxel, a commonly used chemotherapy agent, was kindly gifted from the Oregon State University College of Pharmacy High-Throughput Screening Services Laboratory (HTSSL).^27^

For our free drug analysis in 2D, endometrial cancer cells (HEC-1A, KLE, and Ishikawa) were seeded in 96 well plates with 35 µL of endometrial cancer culture media. After 24 hours of culturing, we performed an eight-point half-log screen ranging from 0.008 to 25 µM. Paclitaxel was diluted with dimethyl sulfoxide (DMSO) and dispensed onto the wells using an automatic liquid handler (D300e Digital Dispenser, Hewlett Packard [H.P.], Corvallis, OR). Media alone and DMSO controls were also evaluated. We first evaluated the IC_50_ values for each cell line using the conventional metric of concentration that reduces the cell viability by 50%. Specifically, we normalized the luminescence or absorbance signal values to the average of the values observed in the absence of the drug, and each replicate was considered as an individual point. The IC_50_ and Hill slope were determined from these results (**Equation 1**). The dose response-inhibition curves were calculated using Prism 8.2.1 software (GraphPad, San Diego, CA).

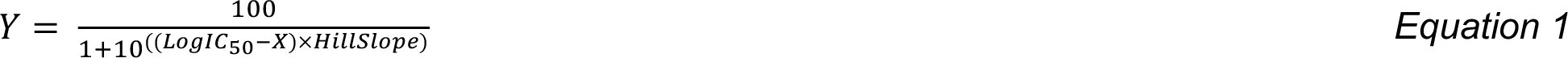

Where Y is the normalized cell response, and X is the log of the drug concentration.

We then evaluated the IC_50_ values in terms of phenotypic responses. Using our optimized hydrogel model, CellTracker blue-labeled endothelial cells and CellTracker green-labeled endometrial cancer cells were seeded in the optimized construct described above and cultured for 24 hours. After 24 hours of culturing, we performed an eight-point half-log screen by dispensing Paclitaxel on top of the 3D *in vitro* models with the automatic liquid handler (D300e, HP) and using DMSO as a vehicle. Paclitaxel loaded in PCL nanoparticles was pipetted by hand into the wells at the same concentration of the free drug. Phenotypic cell responses were measured after 24 and 48 hours of treatment. The IC_50_ response curves were then calculated using the same approach as for cell viability using **Equation 1**.

### Statistical analysis

Statistical analyses were performed using a two-way analysis of variance (ANOVA) with Tukey correction for multiple comparisons or one-way ANOVA when appropriate. Dunnet correction was used in the DOE comparison to account for the significant difference between the predicted hydrogels and Matrigel. All statistical analyses were performed using Prism 8.2.1 software (GraphPad, San Diego, CA); *p* values less than 0.05 were considered statistically significant. Replicates are specified in each experiment, and asterisks denote statistical significance.

## 3. Results

### 3.1 Using design of experiments for developing a 3D in vitro model for endometrial cancer

Using a Design of experiments (DOE) approach, we engineered a hydrogel formulation to support human microvascular endothelial cells (hMVEC) and a hydrogel formulation to support endometrial cancer cells. Using a D-optimal experimental design, seven input variables generated 45 unique hydrogel combinations with a centrally repeated condition. Specifically, we evaluated the influence of the concentration of Col I, fibrinogen and a synthetic polymer PEGDA (10% w/v) or GelMA (7% w/v) in the bottom hydrogel layer; and Col IV, fibronectin, laminin and a PEGDA (10% w/v) or GelMA (7% w/v) in the top hydrogel layer. From the experimental results, we generated multivariant models of microvessel length, endometrial invasion depth, endothelial cell coverage, and endometrial cancer coverage, with predicted vs. observed R^2^ values of 0.89, 0.89, 0.70, and 0.83, respectively, and Q^2^ values ≥0.50 suggesting that the number of model terms adequately fit the model. The R^2^-adjusted values, though slightly lower, remain robust and underline the model’s reliability. Finally, the G-efficiency of the model was 51.32%. Robust model designs have a G-efficiency of 50% or larger ^28^. The ANOVA table showed that one or more input variables significantly influenced each cell type in each hydrogel layer (*p*<0.05), and the interactions were described as linear or two-factor interactions (**Table 1**). These results underscore the significant correlations between the input variables and the measured responses. Endothelial microvessel length ranged from 16 µm to 40 µm and was significantly influenced by the individual concentration of extracellular matrix components and their quadratic terms (**Fig 3A**).

**Table 1.** Hydrogel formulations predicted to maximize or minimize cell phenotypes of interest in the bottom hydrogel (B) or top hydrogel (T). FG = fibrinogen, Syn = synthetic polymer, FN = fibronectin, LMN = laminin.

| Response | [Col I]<sub>B</sub><br>(mg/mL) | [FG]<sub>B</sub><br>(mg/mL) | [Syn]<sub>B</sub> | [Col IV]<sub>T</sub><br>(mg/mL) | [FN]<sub>T</sub><br>(mg/mL) | [LMN]<sub>T</sub><br>(μg/mL) | [Syn]<sub>T</sub> |
| --- | --- | --- | --- | --- | --- | --- | --- |
| Maximize all cell phenotypes of interest | 1.91 | 0.60 | GelMA | 0.13 | 0.17 | 0.50 | PEGDA |
| Minimize all cell phenotypes of interest | 0.70 | 0.70 | PEGDA | 0.03 | 0.16 | 0.65 | PEGDA |

**Figure 3.**
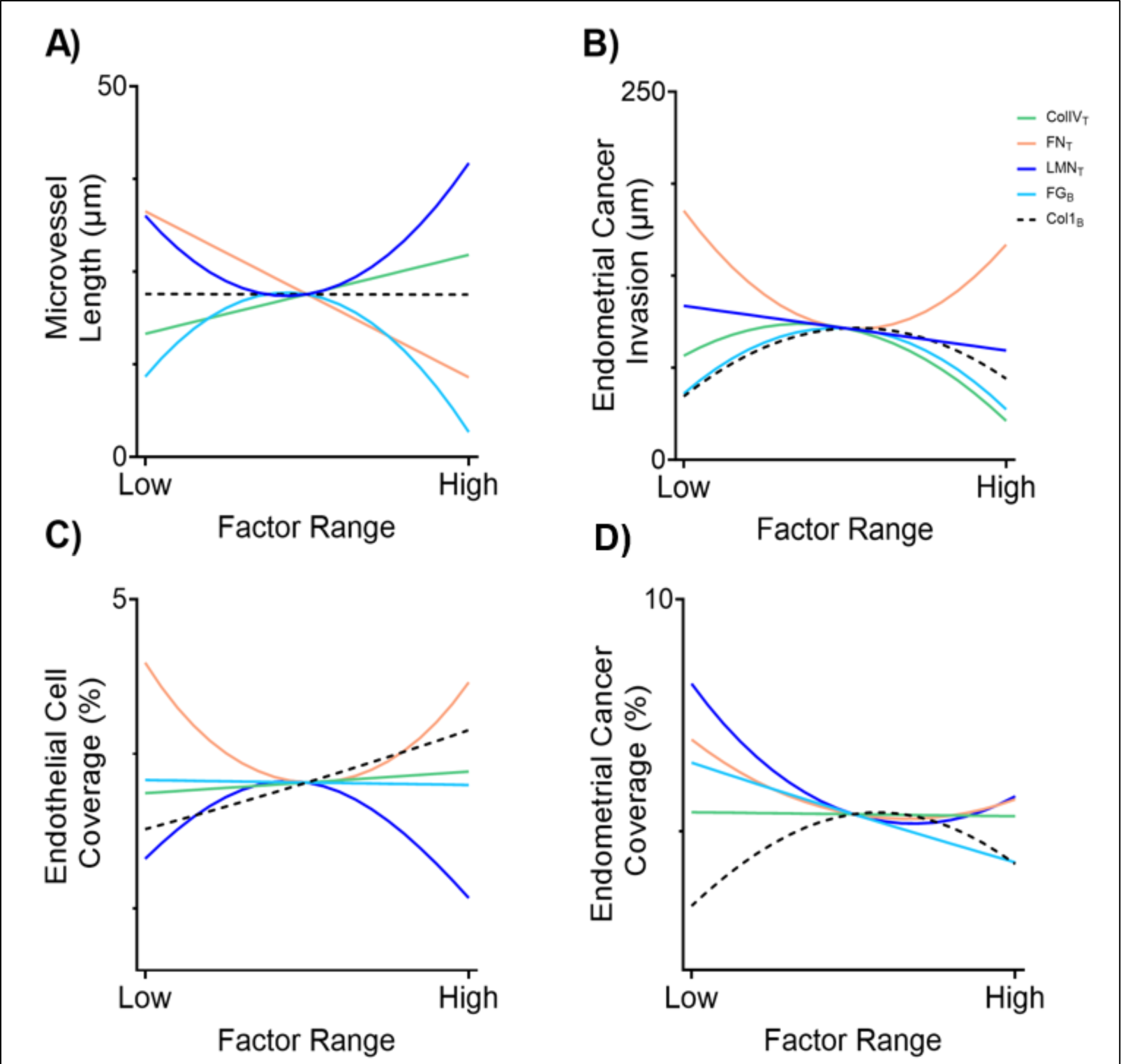
Factor effect plots of the DOE results. Factor effect plots for each cell response as a function of Col1 in the bottom hydrogel ([Col1]B, dashed black line), fibrinogen in the bottom hydrogel ([FG]B, solid teal line), ColIV in the top hydrogel ([ColIV]_T_, solid green line), fibronectin in the top hydrogel (T, solid orange line), and laminin ([LMN]T, solid blue line). (**A**) Microvessel length; (**B**) endometrial cancer invasion; (**C**) endothelial cell coverage; and (**D**) endometrial cancer coverage. Data represented as the predicted cell response to each factor. Data represent factor effect plots generated from predictive equations for each response as a function of the four input variables while other input factors are held constant at their midlevel value. Second order multiple linear regression models for each response formed from N=45 conditions, N=4 per condition.

Specifically, Col IV had a positive linear effect in the bottom hydrogel. The quadratic term did not contribute to microvessel formation. Laminin had a more significant influence on endothelial cell microvessel length when combined with a synthetic polymer and positively influenced microvessel length in a quadratic relationship. These quadratic relationships are better visualized by examining a microvessel length surface response plot as a function of ColIV in the top hydrogel and fibrinogen in the bottom hydrogel (**Fig. 4A**). The concentration of fibrinogen in the bottom hydrogel did not have a significant effect on microvessel length alone. However, it did have a quadratic interaction, which negatively influenced this cell response (**Fig. 3A**). Col I in the bottom layer only showed a significant interaction with PEGDA and GelMA in the bottom layer. All the ECM components had significant interactions with the synthetic polymers in the top and bottom layers. However, PEGDA or GelMA alone and their quadratic interaction did not influence microvessel length.

**Figure 4.**
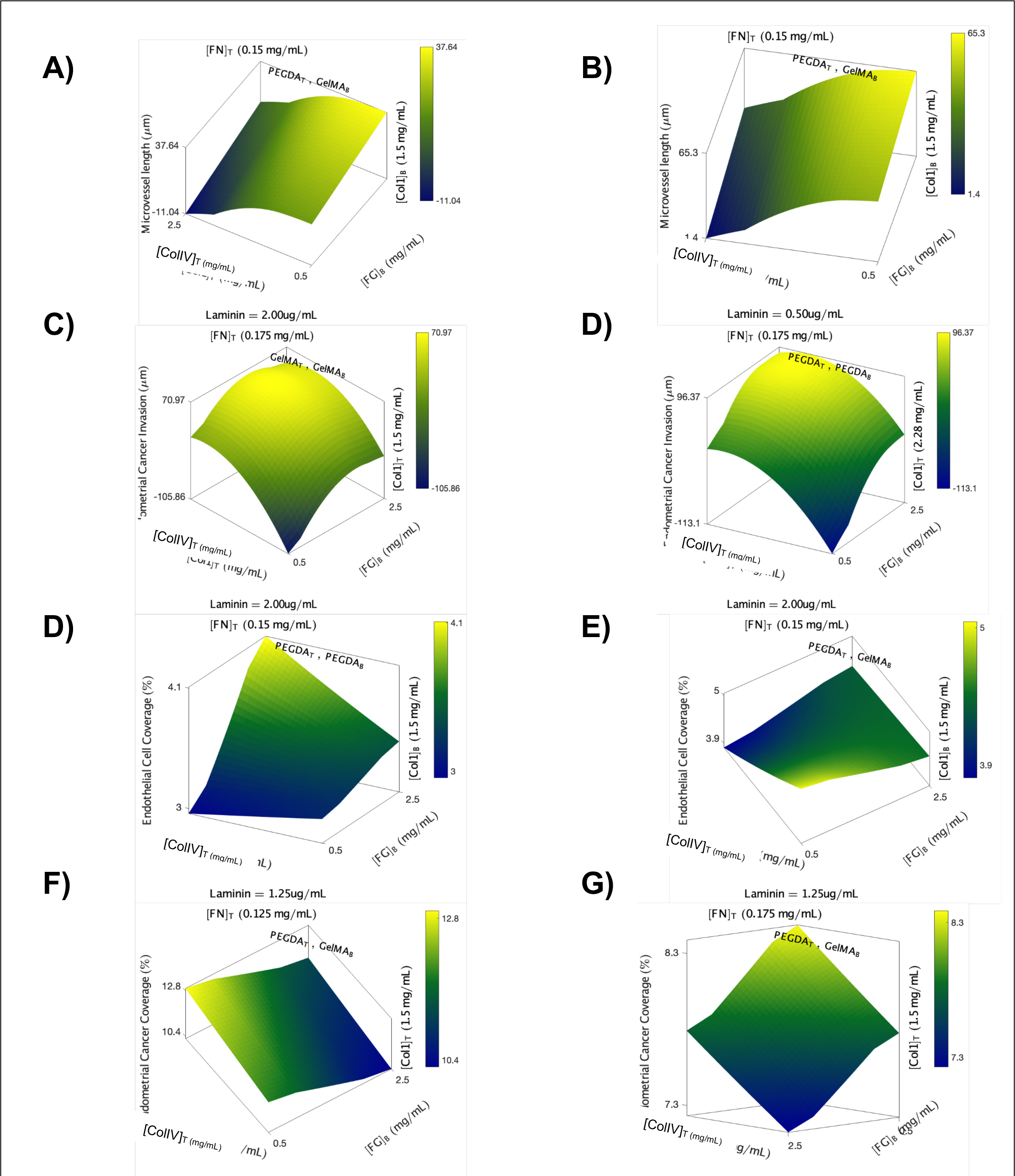
Response surface maps of the DOE results. Response surface maps of hydrogel formulation effects on (**A-B**) Microvessel length (**C-D**) endometrial cancer invasion (**E-F**) endothelial cell coverage (**G-H)** endometrial cancer coverage. Each response surface map demonstrates the predicted output response on the z-axis for a range of ColIV in the top hydrogel ([ColIV]T), Col1 in the bottom hydrogel ([Col1]B), and fibrinogen in the bottom hydrogel ([FG]B) as denoted on the x and y axes at set constant concentrations of fibronectin (T), and laminin in the top hydrogel as denoted at the top and bottom of each graph and set concentrations of the fourth variable ([Col1]T or ([Col1]B) as denoted on the right of each graph. The color bar represents the magnitude of the response from low (dark blue) to high (yellow). Data represent response surface maps generated from predictive equations for each response as a function of the four input variables while other input factors are held constant. Second order multiple linear regression models for each response formed from N=45 conditions, N=4 per condition.

Endometrial cancer invasion, specifically the HEC-1A cell line, ranged from 34 µm to 169 µm and was mostly influenced by the quadratic interactions in the top and bottom hydrogel layers (**Supplemental Table 3**). Col IV, fibrinogen, and Col I had negative quadratic relationships with endometrial cancer invasion, with the maximum cancer cell invasion occurring at the middle concentration range of these ECM components (**Fig. 4C, 3D**). Fibronectin had a positive quadratic influence (**Fig. 3B**). Moreover, it showed only significant interactions with the ECM components in the bottom layer and the synthetic polymers in the top hydrogel layer but not with the synthetic polymers in the bottom layer or Col IV or Laminin (**Table 1**). Laminin was the only component that showed a negative linear relationship with endometrial cancer invasion (**Fig. 3B**). Lastly, the synthetic polymers in the top and bottom hydrogels all had significant interactions with each other (**Fig. 4D**).

Endothelial cell coverage ranged from 3% to 4.2%. Col I in the bottom hydrogel had a positive linear relationship with endothelial cell coverage (**Fig. 3C**). Col IV and fibrinogen in the top hydrogel had a slight yet significant positive influence on endothelial cell coverage. Yet, their individual terms were not significant to this cell response. The quadratic influence of laminin in the top layer negatively affected endothelial cell coverage; meanwhile, the quadratic influence of fibronectin in the top layer positively contributed to endothelial cell coverage (**Fig. 3C**). Switching the synthetic polymers in the bottom layer from PEGDA (**Fig. 4E)** to GelMA **(Fig. 4F)** in the bottom layer increased endothelial cell coverage at low concentrations of fibrinogen in the bottom layer and a mid-range concentration of Col IV in the top layer.

Endometrial cancer coverage ranged from 3% to 8%. All the ECM individual components had a significant effect on cancer coverage except for Col IV in the top hydrogel layer (**Fig. 3D**). Interestingly, laminin and fibronectin in the top layer had a negative quadratic influence on cell coverage (**Fig. 3D**). This was further visualized by examining the surface response curves (**Fig. 4G and 4H**). Fibronectin interacted with the synthetic polymers in the top and bottom hydrogel layers, with a stronger influence in the top hydrogel layer. By increasing fibronectin in the top layer from 0.125 mg/mL to 0.175 mg/mL, the surface response curves showed a maximum endometrial cancer cell coverage at high concentrations of Col IV and fibrinogen and a minimum response at low Col IV and low fibrinogen. The quadratic influence of laminin in cancer coverage showed that by increasing this factor, the coverage decreased from 12.8 % to 8.3% (**Fig. 4H**). Overall, these results showed that endothelial cell response and endometrial cancer response were influenced by the linear and quadratic interactions from the D-Optimal design and opposite cell responses were exhibited when one or two components were changed in the model. Therefore, achieving the ideal hydrogel formulations requires finding a delicate equilibrium between the responses of both cell types.

### 3.2 Validation of DOE Model

The prediction algorithm from MODDE Pro was used to find the hydrogel formulations that would maximize or minimize endothelial cell microvessel length, endometrial cancer invasion, endothelial cell coverage, and endometrial cancer coverage (**Supplemental Table 3**). The design criteria were based on the formulation with a lower probability of failure (less than 10%), and Matrigel was included as a control.

The hydrogel formulation that maximized all phenotypic cell responses (Max All) had a fibrinogen concentration of 0.6 mg/mL, Col I was 1.91 mg/mL, and mixed with GelMA (7% w/v) to form the bottom hydrogel layer. The top layer had a concentration of 0.13 mg/mL Col IV, fibronectin was 0.17 mg/mL, laminin 0.50 μg/mL, and were mixed with PEGDA (10% w/v). This hydrogel had significantly higher microvessel length values compared to other hydrogel formulations (**Fig. 5A**). Furthermore, while Matrigel exhibited a decrease in microvessel length after 12 hours of culturing, the Max All formulation demonstrated stability in microvessel length, maintaining consistent values over 48 hours. This hydrogel also showed a higher endometrial cancer invasion and increasing values over time, compared to other hydrogel formulations, including Matrigel (**Fig. 5B**). The Max all hydrogel formulation showed significant differences between Matrigel and the formulation predicted to minimize all phenotypic cell responses (Min All formulation). Endothelial cell coverage, on average, demonstrated a higher percentage in coverage compared to other hydrogel formulations. Over time, it exhibited a similar trend to other responses (**Fig. 5C**). There were significant differences across all the hydrogel formulations for endothelial cell coverage. Endometrial cancer coverage increased over the 48 hours across all hydrogel formulations, and the Max All formulation displayed a significantly higher cancer coverage compared to the other hydrogel formulations (**Fig. 5D**).

**Figure 5.**
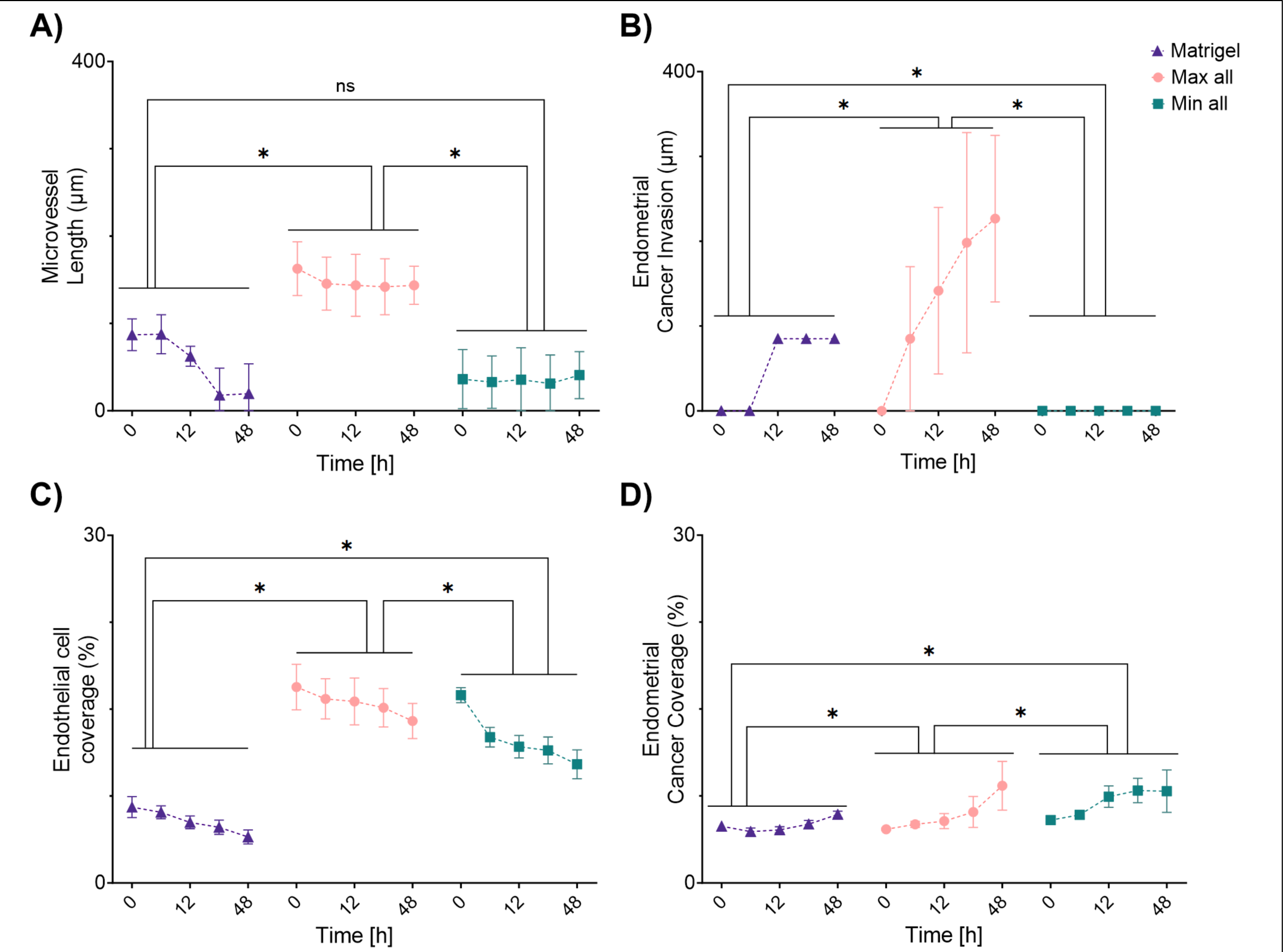
Validation of the DOE predictive models. The desirability function algorithm was used to identify the hydrogel formulations predicted to either maximize or minimize all four of the cell phenotypic behaviors simultaneously (maximize all) and (minimize all). Phenotypic responses, **(A)** microvessel length **(B)** endometrial cancer invasion (**C**) endothelial cell coverage, and (**E**) endometrial cancer coverage were measured every 3 hours for 48 hours. Matrigel was used as a control. * *p* <0.05 compared with the mean of each group. Data were analyzed using a two-way ANOVA with Tukey post-test. Data represent the mean ± SD (n = 4).

The hydrogel formulation designed to minimize all phenotypic responses (Min All) behaved as predicted. Microvessel length and endometrial cancer invasion had the lowest values with this formulation (**Fig. 5A, 5B**). Furthermore, endothelial cell coverage decreased continuously with this hydrogel, maintaining levels lower than those observed with the Max All formulation after 48 hours of culturing (**Fig. 5C**). Regarding endometrial cancer coverage, the Min All hydrogel initially experienced an increase within the first 12 hours. However, following this initial increase, the response remained constant for the subsequent 48 hours, unlike other hydrogel formulations that continually increased cancer coverage throughout the 48-hour experiment (**Fig. 5D**). Due to the limited dynamic range of cell coverage, we concentrated on optimizing microvessel length and cancer invasion in further experiments. We further validated the DOE-predicted model using hydrogel formulations designed to both maximize and minimize microvessel length and endometrial cancer invasion separately (**Fig. 1S**).

### 3.3 Replicating the tumor microenvironment of endometrial cancer

We evaluated the impact of crosstalk between endothelial cells and endometrial cancer cells on microvessel formation and cancer invasion (**Figure 6**). Each cell type was cultured on the optimized DOE hydrogel formulations (Max All) within a co-culture system, where both epithelial and endothelial cell types were present together, and a monoculture system, with only one cell type present. As a control, Matrigel replaced the Max All hydrogel formulation in the top and bottom hydrogel layers. On both Matrigel and the Max All hydrogel formulations, the phenotypic responses were elevated on co-culture compared to monoculture. Specifically, the microvessel length was significantly affected by the hydrogel formulation and cell crosstalk, with co-culture resulting in four times higher microvessel length on the Max All hydrogels compared to Matrigel (**Fig. 6A**). Similarly, endometrial cancer invasion was significantly influenced by hydrogel formulation and cell crosstalk, with three times higher endometrial cancer invasion in the co-culture model with the Max All formulation (**Fig. 6B**).

**Figure 6.**
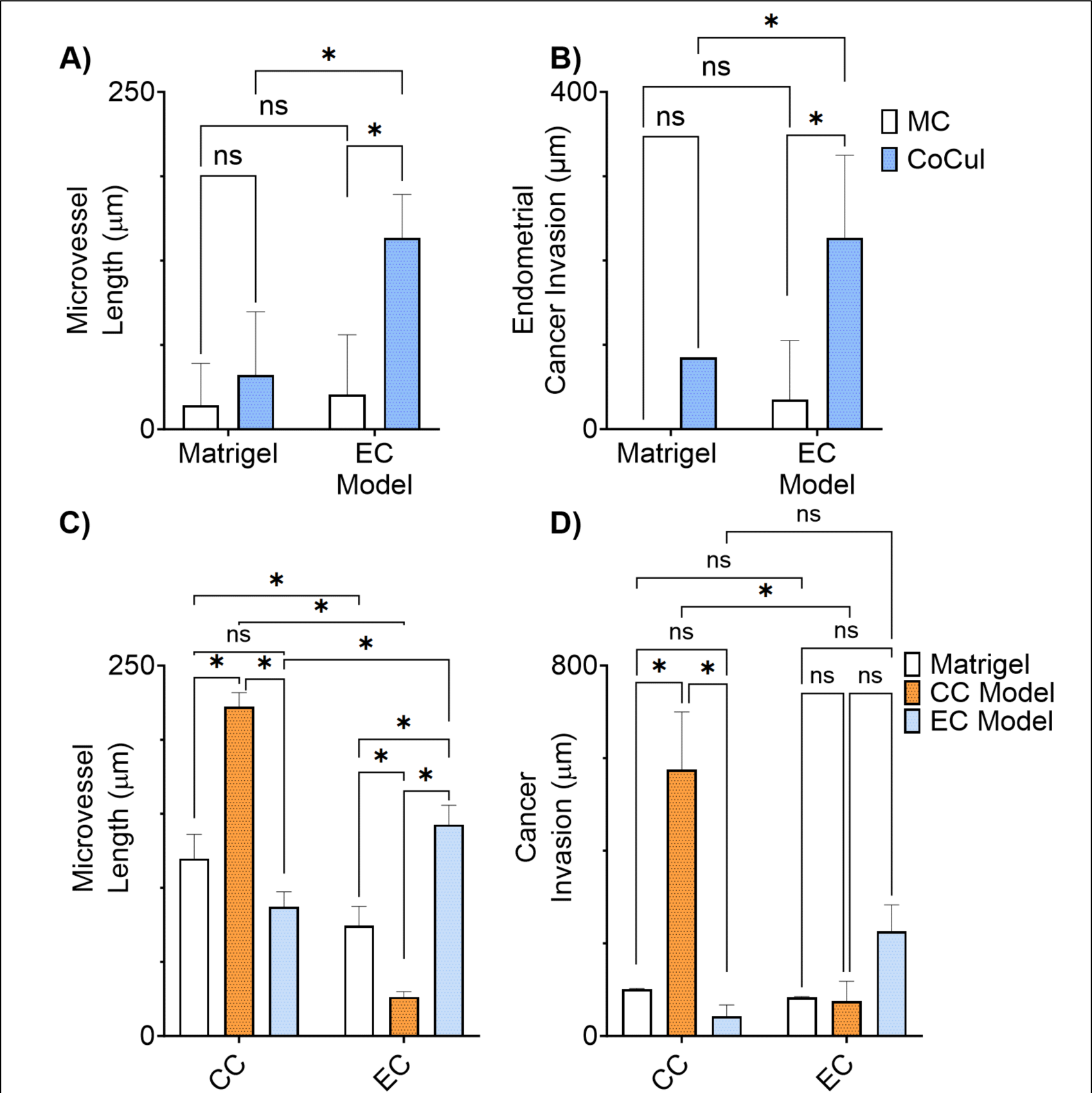
Replicating the tumor microenvironment of endometrial cancer. Cervical cancer cells (HEC-1A) and endothelial cells (hMVEC) were cultured on the “Maximize all” construct in either coculture (CoCul) or monoculture (MC): **(A)** Microvessel length at 12 hours, **(B)** endometrial cancer invasion at 48 hours. Multilayer multicellular models of cervical cancer (C.C.), endometrial cancer (E.C.), and Matrigel were prepared with human microvascular endothelial cells (hMVEC) and either cervical cancer cells (SiHa) or endometrial cancer cells (HEC-1A): (**C**) microvessel length at 12 hours, (**D**) cancer invasion at 48 hours. * *p* <0.05, compared with the mean of each group. Data were analyzed using a Two-way ANOVA, drepresent the mean ± SD (n = 4).

To evaluate the importance of using a specific hydrogel formulation for endometrial cancer, we compared our co-culture Max All hydrogel model (E.C. model), specific to endometrial cancer, with a previously developed hydrogel formulation optimized for cervical cancer phenotypic responses (C.C. model)^19^. Matrigel was used as control. After 48 hours of culturing, noticeable differences in microvessel length were observed when endothelial cells and cancer cells were seeded on their respective matrices. Specifically, culturing cervical cancer cells on the C.C. model and endometrial cancer cells on the E.C. model resulted in a significant increase in microvessel length compared to both the Matrigel control and conditions where cells were not seeded on their specific matrices. Interestingly, when cancer cells were not cultured on their specific matrices, microvessel length decreased even further below the levels observed with Matrigel. Significant differences were present across all the hydrogel formulations (**Fig. 6C**). Similar results were observed in cancer invasion, with C.C. and E.C. cells exhibiting a preference for their specific matrices, showing significantly higher invasion when cultured on their specific matrices. Specifically, cervical cancer invasion significantly decreased when C.C. cells were cultured on top of the E.C. model, with lower cancer invasion than Matrigel. Likewise, endometrial cancer invasion was significantly reduced when E.C. cells were cultured on top of the C.C. model and Matrigel (**Fig. 6D**).

Overall, these results demonstrated the importance of having a specific matrix for each gynecological disease. Moreover, they highlighted the importance of using DOE to design specific hydrogel formulations for each disease that better capture cell response than the current standard material, Matrigel.

### 3.4 Comparative Rheological Analysis of the hydrogels

Similar to our previous work, we compared the viscoelastic properties of our optimized hydrogel formulation (Max All) with Matrigel. Using a rheometer, we measured the elastic modulus (G’) and loss modulus (G“) of the top layer and bottom layer hydrogel formulations from the Max All design. Specifically, our evaluation aimed to discern differences in G’ and G” among Matrigel (9.2 mg/mL), the top hydrogel formulation (1.12 mg/mL ColIV, 0.16 mg/mL fibronectin, 0.05 μg/mL laminin, 10% w/v PEGDA), and the bottom hydrogel formulation (2.5 mg/mL Col1, 2.5 mg/mL fibrinogen, 7% w/v GelMA) (**Fig. 7**). The goal was to identify variations in the viscoelastic properties influencing microvessel formation and endometrial cancer invasion.

**Figure 7.**
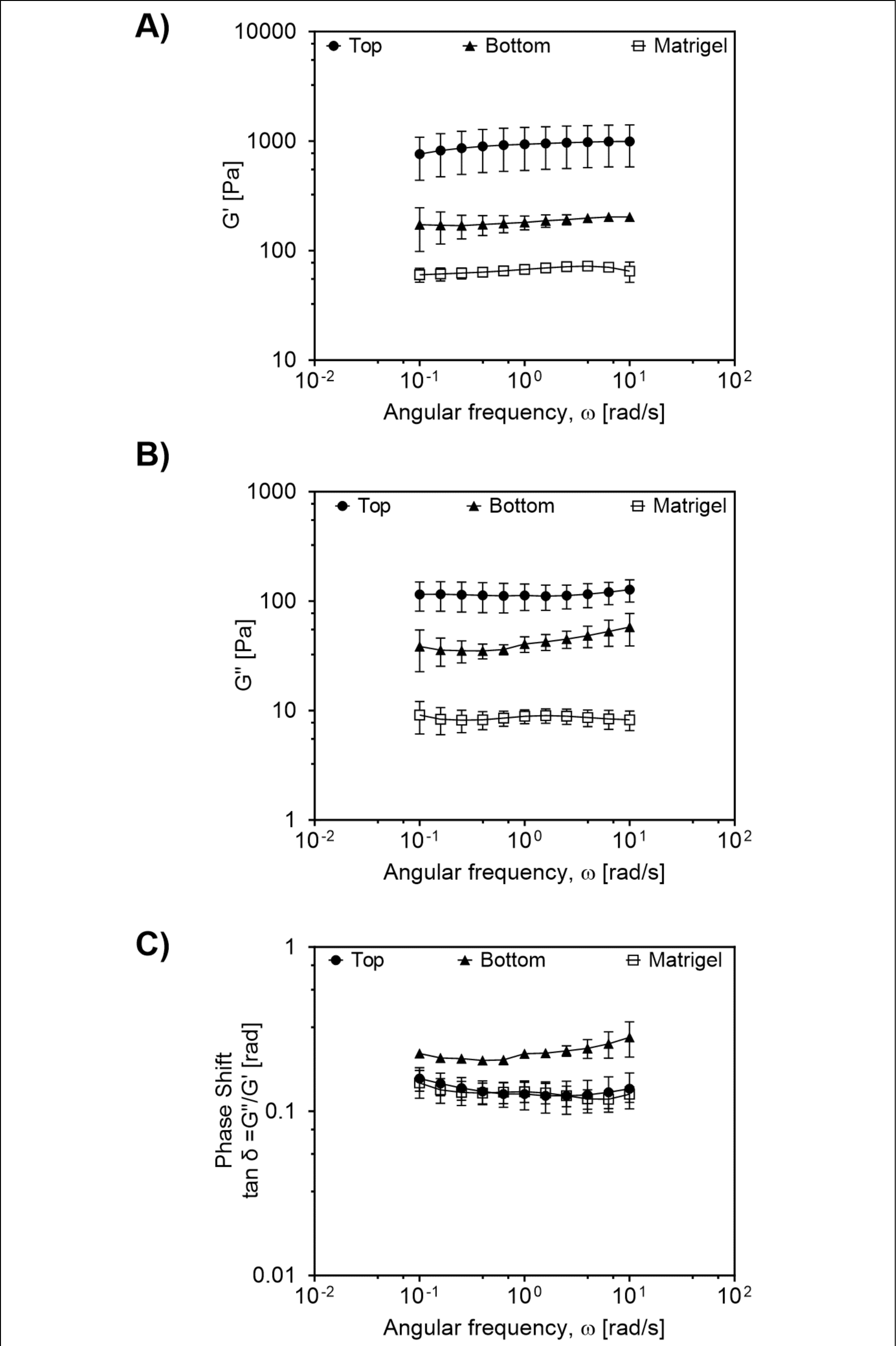
Rheological characterization of hydrogels. **(A)** Storage moduli (G′), (**B**) Loss moduli (G“), and (**C**) Phase shift angle defined as tan δ = G”/G’ for the optimized top hydrogel (Top: 1.12 mg/mL ColIV, 0.16 mg/mL fibronectin, 0.05 μg/mL laminin, 10% w/v PEGDA), the optimized bottom hydrogel (Bottom: 2.5 mg/mL Col1, 2.5 mg/mL fibrinogen, 7% w/v GelMa), and GFD Matrigel (9.2 mg/mL). Frequency sweeps were conducted from 0.1 to 100 rad/sec within the linear viscoelastic (LVE) region at a constant strain amplitude of 5%, and temperature of 37°C. Data represent the mean ± SD (n = 3).

The elastic modulus (G’) is the material’s capacity to store energy during deformation and return to its original shape after stress removal. While the loss modulus (G”) is the material’s capacity to dissipate energy as heat under deformation, indicating its viscous characteristics. In our analysis, the top hydrogel demonstrated significantly higher G’ values than the bottom hydrogel and Matrigel, with Matrigel exhibiting the lowest G’ (**Fig. 7A**). The top hydrogel also had the highest G” value. However, there were no significant differences between the top and bottom hydrogels or Matrigel (**Fig. 7B**). Throughout the frequency sweep analysis, the G’ and G” values of the top hydrogel and Matrigel remained relatively constant. Conversely, the bottom hydrogel layer demonstrated a slight increase in G” towards the end of the analysis. To further understand the viscoelastic behavior of the hydrogels, we evaluated the phase shift angle, represented by tan δ= G”/G’. The top hydrogel exhibited phase shift angles comparable to Matrigel. Conversely, the bottom hydrogel displayed a higher phase shift angle, indicating a more pronounced viscous behavior within this hydrogel. While distinctions were observed between the top, bottom, and Matrigel, all tan δ values were below 1, signifying the retention of viscoelastic properties under shear stress (**Fig. 7C**).

### 3.4 Phenotypic cell response among different endometrial cancer cell lines

To further assess the potential of our optimized multilayer hydrogel model, we conducted a comparative analysis of phenotypic cell responses using three endometrial cancer cell lines seeded on either our Max All formulation or Matrigel. Specifically, hMVEC cells were seeded on the bottom hydrogel layer and co-cultured with either HEC-1A, KLE, or Ishikawa endometrial cancer cell lines seeded on the top layer (**Fig. 8**).

**Figure 8.**
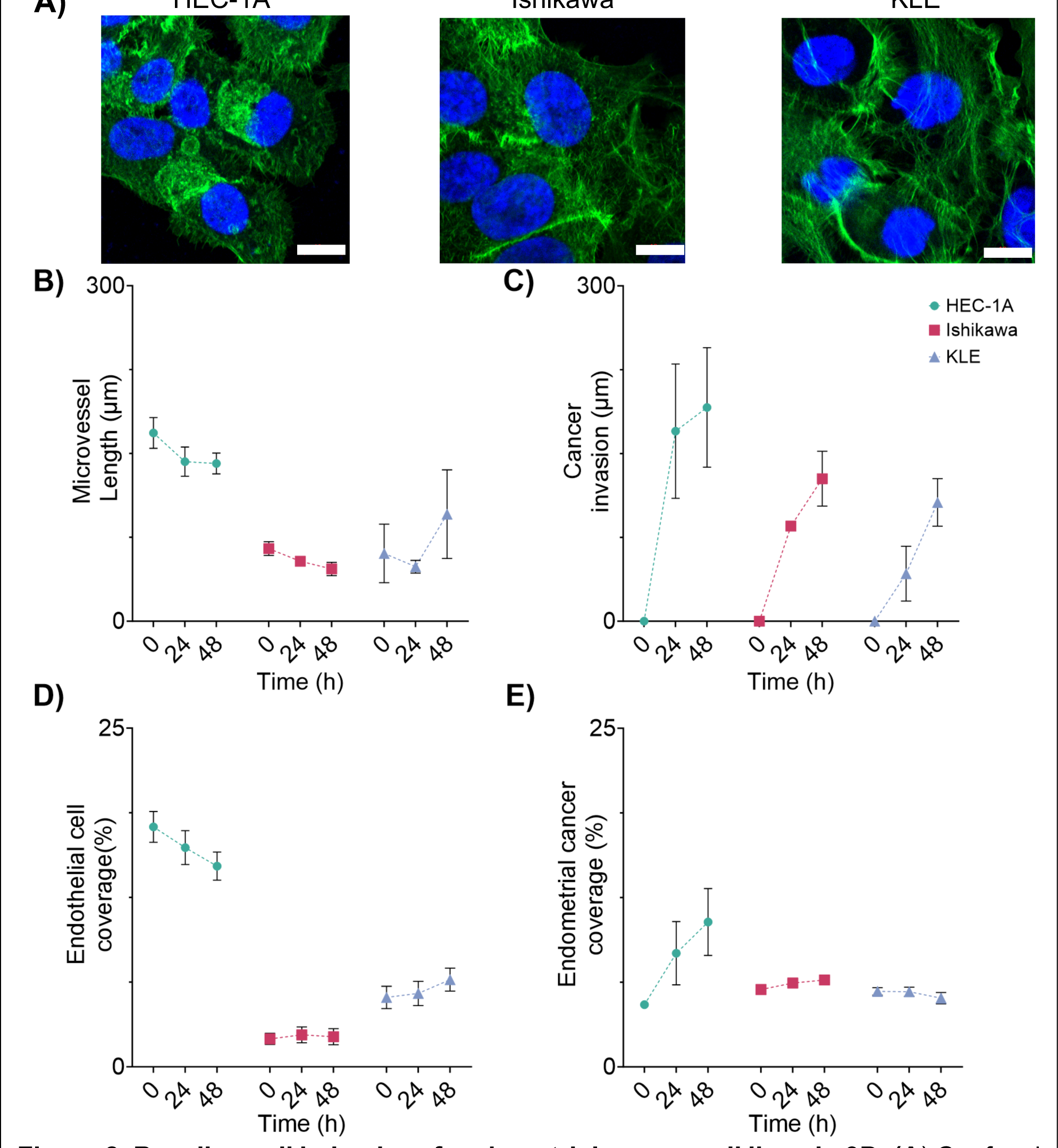
Baseline cell behavior of endometrial cancer cell lines in 3D. (**A**) Confocal images of HEC-1A, Ishikawa and KLE cell lines seeded on TCP and stained with DAPI and phalloidin, scale bar = 10 µm. Endometrial cancer cells were co-cultured with endothelial cells (hMVEC) in 3D using our optimized hydrogel formulation: (**B**) microvessel length (**C**) endometrial cancer invasion (**D**) endothelial cell coverage (**E**) endometrial cancer coverage. Data represent the mean ± SD (n = 4).

Endometrial cancer cells were imaged in the 2D model on poly(L-lysine) coated coverslips (**Fig. 8A**). Microvessel length decreased over time in the experiments with HEC-1A cells and Ishikawa cells. Notably, the construct with KLE cells exhibited a decrease in microvessel length at 24 hours, followed by an increase at the 48-hour mark (**Fig. 8B**). While the constructs with HEC-1A and KLE cells did not show significant differences at 48 hours, distinctions were noted at 0 and 24 hours. Significantly different microvessel formations were observed between the HEC-1A and Ishikawa cells at all time points, and there were no significant differences between constructs with KLE and Ishikawa cells across all time points. Endometrial cancer invasion increased at all time points for all the cell lines (**Fig. 8C**). HEC-1A cells demonstrated the highest level of invasion, followed by Ishikawa cells and KLE cells. There were only significant differences at 24 hours between the constructs with HEC-1A cells and KLE cells. However, after 48 hours of culture, there were no significant differences in invasion depth across all the cell lines.

Endothelial cell coverage showed differences in cell response with all the endometrial cancer cell lines (**Fig. 8D**). Constructs with HEC-1A cells had significantly greater endothelial cell coverage compared to all other endometrial cancer cell lines across all time points, highlighting their unique impact on the endothelial cell environment. Ishikawa cells exhibit a slight increase in coverage at 24 hours, followed by a decrease at 48 hours. In contrast, KLE cells displayed a consistent increase in endothelial cell coverage over time. Endometrial cancer coverage showed contrasting trends across the various cell lines within the constructs compared to endothelial cell coverage. Constructs containing HEC-1A and Ishikawa cells increased cancer cell coverage over time, diverging from the endothelial cell behavior. In contrast, the constructs with KLE cells demonstrated a reduction in cancer cell coverage after the initial 24 hours (**Fig. 8E**). There were only significant differences at the 48-hour mark between the models with HEC-1A cells and the other cell lines. However, models containing KLE and Ishikawa cells showed no significant differences at 48 hours. These findings underscore the model’s dynamic capacity to facilitate diverse responses among endometrial cancer cell lines in their interaction with endothelial cells and the ECM components within the hydrogel formulations. Furthermore, results showed differences in endometrial cancer response over time and endothelial cell response with cancer cells from different tumor sites.

### 3.5 Comparative dose-response analysis of Paclitaxel

We conducted a comparative analysis of the response of three endometrial cancer cell lines to Paclitaxel, a widely used chemotherapy drug. Our investigation encompassed these cells cultured in both traditional tissue culture plastic (2D) and within our specialized multilayer hydrogel model (3D). Our focus was on assessing the impact of Paclitaxel on cell viability and phenotypic responses after drug exposure at 24 hours (**Fig. 2S**) and 48 hours (**Fig. 9**) through an eight-point half log dose-response assessment. Overall, for all cell lines the dose response-inhibition curves demonstrated increased fit at 48 versus 24 hours.

**Figure 9.**
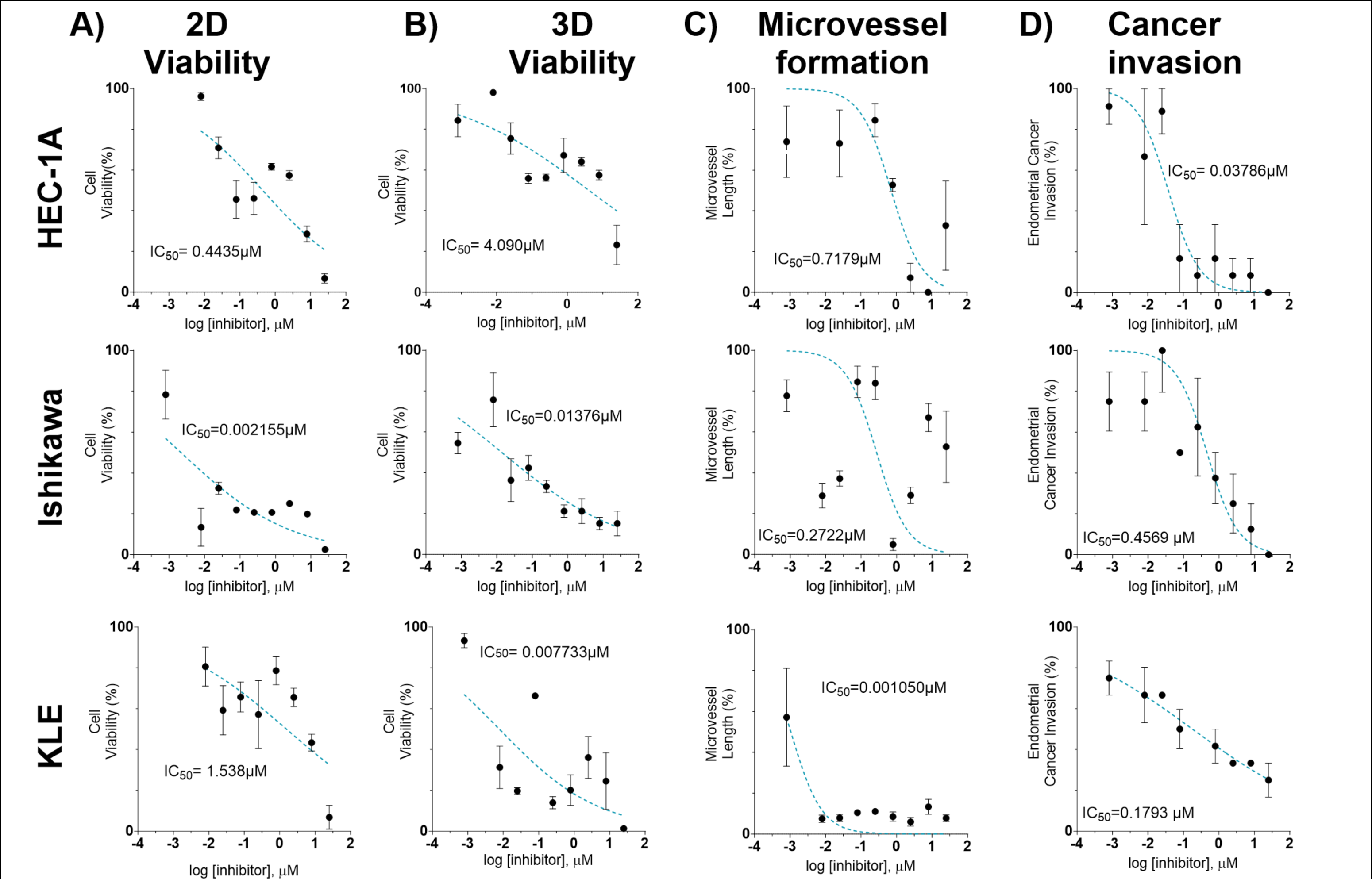
*In vitro* dose response curves of human endometrial cancer cell lines treated with paclitaxel as free drug for 48 hours. Cancer cells were seeded in 96 well plates either on TCP or co-cultured with endothelial cells (hMVEC) in 3D using our optimized hydrogel formulation. Cells were treated with 0.008 – 25 µM of paclitaxel and cell response was evaluated at 48 hours post dosage. Each cell response is normalized to the average of the values observed in the absence of drug. Cell viability and phenotypic cell response were measured in **(A)** HEC-1A **(B)** Ishikawa, and (**C**) KLE cells. Data represent the mean ± SD (n = 4).

**Table 3.**
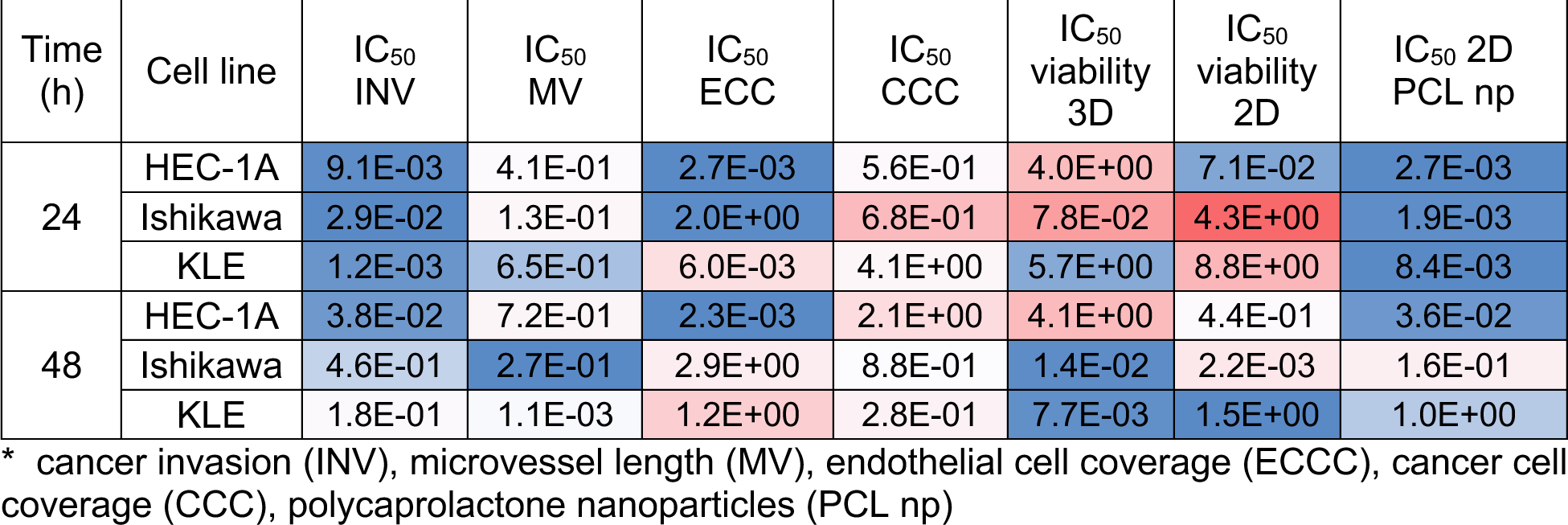
Summary of IC_50_ values for the endometrial cancer cell lines after 24 hours and 48 hours of exposure to Paclitaxel. The drug was delivered as free and encapsulated in PCL nanoparticles on top of our 2D and 3D *in vitro* models.

Cell viability in the 2D models showed differences in the IC_50_ values across the cell lines after 48 hours of culturing (**Fig. 9A**). The highest IC_50_ value was observed in the KLE cells, and the lowest IC_50_ value was observed in Ishikawa cells. Cell viability in the 3D models exhibited a different trend, revealing higher resistance to Paclitaxel in HEC-1A and Ishikawa cells compared to KLE. Additionally, the IC_50_ response curves were better fitting in the 3D models than the 2D models at both 48 hours and 24 hours (**Fig. 9B**, **Fig. 2S**). At 48 hours, the HEC-1A cells exhibited a higher IC_50_ value for microvessel length to Ishikawa and KLE cells (**Fig. 9C**). Conversely, Ishikawa cell models demonstrated a less defined fit in the dose-response curve for microvessel formation, displaying similar IC_50_ values at both 24 and 48 hours (**Fig. 2S**). In contrast, the KLE cell models depicted a consistent dose-response curve in microvessel formation, aligning closely with cell viability in 3D at the 48-hour mark (**Fig. 9C**).

Endometrial cancer invasion demonstrated the best fit dose-response curves across all evaluated metrics (**Fig. 9D**). Among the cell lines, the Paclitaxel-insensitive HEC-1A cells demonstrated the lowest IC_50_ value in cancer invasion across cell lines. In contrast, the Paclitaxel-insensitive Ishikawa cells displayed the highest IC_50_ value compared to the other cancer cell lines and to both 2D and 3D cell viability results. Conversely, the Paclitaxel-sensitive KLE cells exhibited a more favorable fit in the dose-response curve for endometrial cancer invasion in comparison to cell viability and microvessel formation metrics (**Fig. 9D)**. After 24 hours of drug exposure, endometrial cancer cells HEC-1A and KLE still showed Paclitaxel resistance, reflected in the poor fit of the dose-response curve for cancer invasion, a response that notably improved after 48 hours of exposure (**Fig. 2S**). This progression highlights the temporal evolution of their response to Paclitaxel in the context of invasion dynamics (**Fig. 2S**). Endothelial cell coverage and endometrial cancer coverage were also evaluated with Paclitaxel after 24 hours and 48 hours of dosage (**Fig. 3S**). However, the metrics did not exhibit consistent dose-response curves.

We then evaluated the IC_50_ values of Paclitaxel loaded PCL nanoparticles on the three endometrial cancer cell lines. Similar to the free drug, the dose-response curves at 48 hours displayed improved fitting compared to the 24-hour curves (**Fig. 4S**). At both time points, KLE exhibited the highest IC_50_ value, similar to 2D viability with free drug. At 48 hours, HEC-1A cells exhibited increased chemo-resistance when treated with the free drug compared with the PCL nanoparticles. However, Ishikawa cells demonstrated a higher IC_50_ indicative of less chemo-resistance when exposed to Paclitaxel encapsulated in PCL nanoparticles. Moreover, the Paclitaxel-sensitive KLE cells displayed a higher IC_50_ value at 48 hours with both dosing methods compared with the free drug.

These results showed that our 3D *in vitro* platform could capture differences in cell response and viability after being exposed to Paclitaxel. Using the phenotypic cell response elucidated specific behavioral characteristics not discernible through conventional cell viability assays. These findings highlight the capacity of our model to provide a more comprehensive understanding of cellular reactions to Paclitaxel, including comparing between traditional free drug and novel drug-loaded nanoparticles, shedding light on nuanced responses that extend beyond traditional viability measures.

### 3.6 Comparison of cell response to Paclitaxel delivered as a free-drug vs. encapsulated in PCL nanoparticles in 3D in vitro model

In our optimized 3D multilayer *in vitro* model, we conducted a comparative study analyzing cell viability and phenotypic cell responses with Paclitaxel, administered as both a free drug and encapsulated in PCL nanoparticles. Endothelial and endometrial cancer cells were seeded into the 3D model and cultured for 24 hours before treatment. Based on the earlier experiments, the highest IC_50_ value for each cell line at all time points was chosen (**Table 2**), determining the Paclitaxel concentration applied atop the constructs.

**Table 2.** ANOVA table from the DOE models. ANOVA table for the DOE model and interplay between input and output variables with their corresponding *p-*value. R^2^ = coefficient of correlation. Q^2^ =model predictive power. Confidence level 95%. Non-significant values are stated <u>with a slash (-)</u>

| Response | Endothelial microvessel length | Endometrial cancer invasion | Endothelial cell coverage | Endometrial cancer coverage |
| --- | --- | --- | --- | --- |
| p-values |  |  |  |  |
| model | <0.0001 | <0.00001 | <0.0001 | <0.0001 |
| Col1<sub>B</sub> | - | - | 0.0015 | 0.024 |
| FG<sub>B</sub> | - | - | - | 0.000022 |
| Col IV<sub>T</sub> | 0.0107 | 0.0017 | - | - |
| FN<sub>T</sub> | 0.00001 | - | - | 0.0072 |
| LMN<sub>T</sub> | - | 0.0214 | - | 0.000012 |
| Syn<sub>T</sub> (PEGDA) | - | 0.0346 | 0.0044 | - |
| Syn<sub>T</sub> (GelMA) | - | 0.0346 | 0.0044 | - |
| Syn<sub>B</sub> (PEGDA) | - | - | 0.0001 | 0.0120 |
| Syn<sub>B</sub> (GelMA) | - | - | 0.0001 | 0.0120 |
| Col1<sub>B</sub><sup>2</sup> |  | 0.0153 |  | 0.0046 |
| FG<sub>B</sub><sup>2</sup> | 0.0080 | 0.0054 |  |  |
| ColIV<sub>T</sub><sup>2</sup> |  | 0.0164 |  |  |
| FN<sub>T</sub><sup>2</sup> |  | 0.0012 | 0.0128 | - |
| LMN<sub>T</sub><sup>2</sup> | 0.0067 |  | 0.0054 | 0.0098 |
| ColIV<sub>T</sub> * Col1<sub>B</sub> |  | - | 0.0150 |  |
| ColIV<sub>T</sub> * FG<sub>B</sub> |  | - | 0.0245 |  |
| ColIV<sub>T</sub> * LMN<sub>T</sub> |  | - |  | 0.0133 |
| FN<sub>T</sub> * FG<sub>B</sub> |  | 0.0001 |  |  |
| FN<sub>T</sub> * Col1<sub>B</sub> | 0.00002 | 0.0110 |  |  |
| FN<sub>T</sub> * LMN<sub>T</sub> | 0.0025 |  |  |  |
| FG<sub>B</sub> * LMN<sub>T</sub> | 0.0271 | 0.0019 |  | - |
| Col1<sub>B</sub> * LMN<sub>T</sub> |  | 0.0180 |  |  |
| FG<sub>B</sub> * Syn<sub>T</sub> (PEGDA) | 0.00001 |  | 0.0336 |  |
| FG<sub>B</sub> * Syn<sub>T</sub> (GelMA) | 0.00001 |  | 0.0336 |  |
| FG<sub>B</sub> * Syn<sub>B</sub> (PEGDA) | 0.0001 | 0.0290 | - |  |
| FG<sub>B</sub> * Syn<sub>B</sub> (GelMA) | 0.0001 | 0.0290 | - |  |
| Col1<sub>B</sub> * Syn<sub>B</sub> (PEGDA) | 0.0203 |  |  | - |
| Col1<sub>B</sub> * Syn<sub>B</sub> (GelMA) | 0.0203 |  |  | - |
| Col1<sub>B</sub> * Syn<sub>T</sub> (PEGDA) | - | - |  | 0.0306 |
| Col1<sub>B</sub> * Syn<sub>T</sub> (GelMA) | - | - |  | 0.0306 |
| ColIV<sub>T</sub> * Syn<sub>T</sub> (PEGDA) |  | 0.0362 |  |  |
| ColIV<sub>T</sub> * Syn<sub>T</sub> (GelMA) |  | 0.0362 |  |  |
| ColIV<sub>T</sub> * Syn<sub>B</sub> (PEGDA) | 0.0452 |  | - |  |
| ColIV<sub>T</sub> * Syn<sub>B</sub> (GelMA) | 0.0452 |  | - |  |
| FN<sub>T</sub> * Syn<sub>B</sub> (PEGDA) |  | - | - | 0.0090 |
| FN<sub>T</sub> * Syn<sub>B</sub> (GelMA) |  | - | - | 0.0090 |
| FN<sub>T</sub> * Syn<sub>T</sub> (PEGDA) | 0.0002 | 0.0006 |  | 0.0018 |
| FN<sub>T</sub> * Syn<sub>T</sub> (GelMA) | 0.0002 | 0.0006 |  | 0.0018 |
| LMN<sub>T</sub> * Syn<sub>T</sub> (PEGDA) |  | 0.0023 |  |  |
| LMN<sub>T</sub> * Syn<sub>T</sub> (GelMA) |  | 0.0023 |  |  |
| LMN<sub>T</sub> * Syn<sub>B</sub> (PEGDA) | 0.0000002 |  |  |  |
| LMN<sub>T</sub> * Syn<sub>B</sub> (GelMA) | 0.0000002 |  |  |  |
| Syn<sub>T</sub> (PEGDA) * Syn<sub>B</sub> (PEGDA) |  | 0.0061 |  |  |
| Syn<sub>T</sub> (PEGDA) * Syn<sub>B</sub> (GelMA) |  | 0.0061 |  |  |
| Syn<sub>T</sub> (GelMA) * Syn<sub>B</sub> (PEGDA) |  | 0.0061 |  |  |
| Syn<sub>T</sub> (GelMA) * Syn<sub>B</sub> (GelMA) |  | 0.0061 |  |  |
| Lack of fit | 0.111 | 0.682 | 0.017 | 0.004 |
| Validation metrics |  |  |  |  |
| Total sample size (N) | 44 | 43 | 45 | 45 |
| Degree of Freedom (DF) | 24 | 17 | 29 | 28 |
| R<sup>2</sup> | 0.89 | 0.89 | 0.70 | 0.83 |
| R<sup>2</sup>-adjusted | 0.81 | 0.72 | 0.6 | 0.74 |
| Q<sup>2</sup> | 0.58 | 0.39 | 0.27 | 0.56 |

Constructs with HEC-1A cells were treated with 4.02 µM of Paclitaxel, either as a free drug or loaded in PCL nanoparticles. Both delivery methods resulted in decreased viability and phenotypic responses. Paclitaxel, dispensed as a free drug was more effective in reducing endometrial cancer invasion. There were no significant differences between the dosing methods; however, both methods showed reduced cell viability and phenotypic cell responses compared to the control group (**Fig. 10A**).

**Figure 10.**
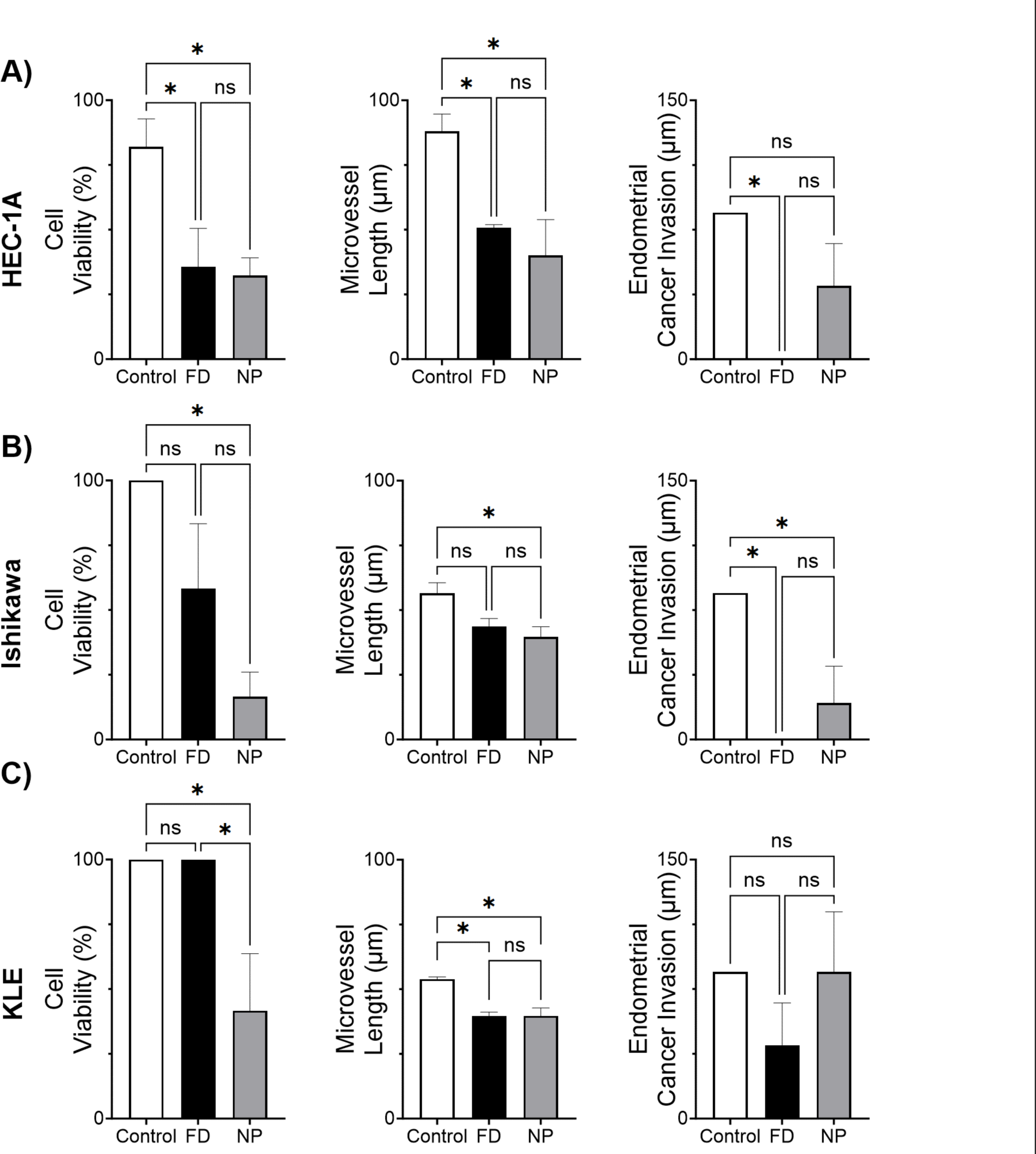
*In vitro* response of human endometrial cancer cell lines in 3D treated with paclitaxel as either free drug or encapsulated in PCL. Endometrial cancer cells were co-cultured with endothelial cells (hMVEC) in 3D using our optimized hydrogel formulation. Cells were treated with paclitaxel as either free drug (FD), or loaded in PCL nanoparticles (NP) for 48 hours, at which point cell response was evaluated. Each cell response is normalized to the average of the values observed in the absence of drug. Cell viability and phenotypic cell response were measured in **(A)** HEC-1A **(B)** Ishikawa, and (**C**) KLE cells. * *p* <0.05 compared with the mean of each group. Data were analyzed using a one-way ANOVA with Tukey post-test. Data represent the mean ± SD (n = 4).

3D cultures with Ishiwaka cells were dosed with Paclitaxel at a concentration of 4.34 µM for both free drug and PCL nanoparticles. Similar to constructs with HEC-1A cells, there was a reduction in cell viability with the PCL nanoparticles, and invasion was decreased with the free drug. Again, there were no significant differences between free drug and PCL nanoparticles. However, only the PCL nanoparticles showed significant differences in the reduction of cell viability compared to the control group (**Fig. 10B**).

The KLE cell models were treated with Paclitaxel at 8.78 µM, revealing the highest resistance among all the evaluated cancer cell lines in both cell viability and endometrial cancer invasion (**Fig. 10C**). Notably, significant differences emerged between the two dosing methods in terms of cell viability. Specifically, only the PCL nanoparticles effectively reduced cell viability, exhibiting significant differences compared to the control group. In contrast, both treatments resulted in a similar decrease in microvessel length, with significant differences only observed compared to the control group. Moreover, the impact on endometrial cancer invasion was solely observed in response to the free drug.

## 4. Discussion

We developed a unique multilayer multicellular model designed to replicate the complex architecture of the endometrium. Comprising two specifically formulated hydrogel layers, our model supports both endothelial and endometrial cacer cells, facilitating cross-talk that mirrors intercactions within the tumor microenvironment. This innovative model enabled us to explore dynamic responses of different endometrial cancer cell lines to Paclitaxel in either its free form or encapsulated in novel PCL nanoparticles, revealing nuanced cell behaviors not captured by convential 2D cell viability assays. Additionally, our model’s cell phenotype analysis in a 96-well plate format positions it as a seamless fit for high-throughput screening methods.

Using a Design of Experiments (DOE) approach, we developed two hydrogel formulatiopns to maximize endometrial cancer invasion and microvessel formation. The two distinct formulations for the top and bottom layers incorporated a blend of ECM components and synthetic polymers, leading to enhanced metastatic behaviors compared to the traditional model of Matrigel.

Specifically, the bottom hydrogel layer was developed using a blend of Col I, fibrinogen, and synthetic polymers—GelMA or PEGDA. Echoing our previous findings,^19^ we found that varying Col I and fibrinogen concentrations within the range of 0.5 to 2.5 mg/mL significantly influenced microvessel formation and cancer invasion. Specifically, 1.91 mg/mL Col I and 0.6 mg/mL fibrinogen maximized microvessel length and endometrial cancer invasion, demonstrating significant interactions with each other. This aligns with previous studies that have evaluated the influence of Col I in promoting microvessel formation^29^ and its influence on microvessel formation when combined with fibrin hydrogels.^30^ Following a similar approach for the bottom hydrogel layer, the top hydrogel layer was developed using a combination of Col IV, fibronectin, laminin, and either PEGDA or GelMA. These components were chosen based on their influence on endometrial cancer behavior.^17,31,32^ We determined that a concentration of 0.13 mg/mL Col IV maximized endometrial cancer invasion, falling within the mid-range of our tested spectrum (0.01 to 0.2 mg/mL). Col IV is a predominant structural component of cellular basement membranes and has been demonstrated to have a complex, context-dependent relationship with endometrial cancer progression.^33,34^ Furthermore, we found that 0.5 mg/mL laminin maximized cancer invasion and significantly affected microvessel length and endothelial cell coverage, which was at the bottom of our tested spectrum (0.5 to 2 mg/mL). Interestingly, compared to the concentrations in Matrigel (approximately 2.8 mg/mL Col IV and 5.5 mg/mL laminin), our optimal levels were substantially lower, highlighting potential reasons for Matrigel’s limitations in promoting endometrial cancer metastatic behaviors. In addition to Col IV and laminin, fibronectin is a major component of the endometrial cancer tumor microenvironment and has been previously demonstrated to promote endometrial cancer growth and invasion.^1,17,35^ We evaluated fibronectin from 0.125 to 0.175 mg/mL and found that the concentration that maximized endothelial and endometrial cancer metastatic behavrior was 0.17 mg/mL, at the higher end of our tested range. All three of these ECM components had significant interactions with each other as well as ECM components in the bottom hydrogel, again highlighting the importance of capturing the layered architecture and multicellular nature of the endometrial cancer tumor microenvironment. Lastly, the incorporation of synthetic polymers, PEGDA and GelMA, in both hydrogel layers was found to have significant interactions with the ECM components and significantly affect cell behavior. Optimal results were achieved with PEGDA in the top layer and GelMA in the bottom layer. This study is the first to report a combination of GelMA, Col I, and fibrinogen to support endothelial cells as well as the first to report a combination of PEGDA, Col IV, laminin, and fibronectin to support endometrial cancer cells. The addition of these synthetic polymers to the ECM proteins allowed flexibility of the mechanical properties, supporting the co-culture system.

To investigate the differences in material properties between our optimized hydrogel formulations and Matrigel, we evaluated their visoelastic properties. The top hydrogel formulation had the highest viscoelasticity and mechanical strength with a storage modulus (G’) of 916 Pa and loss moduluis (G“) of 115 Pa. This is in contrast to the bottom hydrogel (148 Pa G’, 43 Pa G’’), formulation and Matrigel (66 Pa G’, 9 Pa G’’). Importantly, the viscoelastic properties of the top hydrogel closely align with physiological conditions; the elastic modulus of a nonpregnant endometrium is approximately 250 Pa and the complex modulus of uterine tissue is approximately 100 Pa.^36,37^ Hence, our top hydrogel more accurately replicates the stiffness in native endometrium compared to Matrigel which is significantly softer. This increased stiffness in our hydrogel is attributed to the incorporation of PEGDA and collagen 1, providing the mechanical rididity that Matrigel lacks. This similarity in mechanical properties to those of endometrial tissue likely contributes to the enhanced endometrial cancer invasion observed in our model, as clinical data has demonstrated a correlation between increased stiffness and endometrial cancer progression.^38^

To validate our model in different stages of endometrial cancer, we evaluated three endometrial cancer cell lines: HEC-1A (grade 2 endometrial carcinoma), Ishikawa (stage 2, endometrial adenocarcinoma) and KLE (metastatic adenocarcinoma).^2^ Consistent with previous studies, HEC-1A cells were the most invasive and our model provided new insights into how these cell lines interacted with endothelial cells within a 3D environment. When treated with Paclitaxel, our 3D platform reveled significant differences in drug sensitivities between the three cell lines in terms of viability that were in contrast to what was observed in 2D. Although the KLE cells had the highest IC_50_ values in 2D, indicating reduced chemosensitivity, the HEC-1A cells had the highest IC_50_ values in 3D. This may be due to increased proliferation of KLE cells in monoculture following exposure to chemotherapy drugs.^4^ HEC-1A cells are known to be paclitaxel insentitive while KLE cells are known to be sensitive to paclitaxel.^23^ In our studies, the 2D IC_50_ values do not support the known Paclitaxel sensitivity of the cell lines, however, the 3D models align with previous findings which show the 3D model better predicts the *in vivo* behavior.^39^

In addition to cell viability, we assessed cancer invasion and microvessel length, to provide a broader evaluation of metastatic behavior. Interestingly, our findings revealed that Paclitaxel treatment had a more pronounced effect on KLE invasion compared to KLE cell viability. Expanding on this notion and exploring phenotypic cell responses in other cancer cells, we observed distinctive patterns for Ishikawa and HEC-1A cells. Models featuring HEC-1A cells exhibited increased resistance in microvessel formation, suggesting a heightened resilience to the drug. Conversely, Ishikawa cells displayed elevated resistance in cancer invasion compared to the other cell lines, implying a unique response profile. These nuanced findings underscore the importance of dissecting specific phenotypic responses, as different endometrial cancer cell lines demonstrated varied susceptibilities to Paclitaxel in the 3D microenvironment.

Paclitaxel is one of the most commonly used chemotherapy agent for treating endometrial cancer. However, it is traditionally administered intravenously and its effectiveness can be limited by systemic toxicity and poor solubility.^4040^ Addressing these issues, we developed a novel poly(caprolactone) (PCL) nanoparticle delivery system for Paclitaxel. This approach is part of a broader trend in cancer treatment, where nanoparticle-based delivery systems are increasingly being investigated.^40^ For example, nanoparticle albumin-bound Paclitaxel (nab-Paclitaxel) was the first nanotechnology-based cancer treatment and is clinically approved for the treatment of metastatic breast and pancreatic cancer.^41,42^ Our nanoparticle delivery system differs by encapsulating the paclitaxel with in PCL nanoparticles. Thus, our Paclitaxel-loaded PCL nanoparticles create a treatment that overcomes the poor solubility issue associated with free paclitaxel. By encapsulating the paclitaxel in PCL, the paclitaxel can then slowly diffuse through the PCL layer into the tumor environment where the particles collect due to the EPR effect. In turn, this helps to reduce the systemic toxicity. Furthermore, PCL particles can also increase the sensitivity of the cancer cell to the paclitaxel by the nanoparticles being endocytosed into the cell and the paclitaxel diffusing out into the cell, resulting in lower required doses to observe the same therapeutic effect.

Finally, we then used our 3D model to evaluate the efficacy of our PCL nanoparticle delivery system. By using our 3D model we gained insights beyond cell viability, enabling us to evaluate the effects on endometrial cancer invasion and microvessel formation over time. Furthermore, as our 3D model was developed in a 96-well plate format, we were able to use automatic dispensing equipment to perform our dose-response studies. The administration of Paclitaxel, both free and nanoparticle-encapsulated, revealed distinct responses across different endometrial cancer cell lines. Overall, the PCL nanoparticles significantly reduced cell viability in all three cell lines, aligning with previous work demonstrating the effectiveness of polymeric nanoparticles as drug delivery systems.^2,43,44^ Importantly, here we show for the first time that by using Pacliaxel-loaded PCL nanoparticles we can induce chemosensitivity in a both a Pacliaxel-sensitive endometrial cancer cell line (KLE) and a classically Pacliaxel-insensitive endometrial cancer cell lines (Hec-1A and Ishikawa). Notably, decreased cell viability did not correlate with decreased endometrial cancer invasion or decreased microvessel length. While the Pacliaxel-loaded nanoparticles reduced endometrial cancer invasion and microvessel formation compared to the untreated controls, there were no significant differences between the free drug and the nanoparticles. These data highlight the importance of evaluating multiple facets of endometrial cancer metastasis beyond cell viability. Furthermore, this methodology aligns with the FDA’s recent recognition of the value of 3D models in preclinical data for drug development.^45^ By incorporating these advanced 3D models, we can contribute to more relevant and reliable preclinical data, potentially accelerating the development and approval of new cancer treatments. This integration of novel drug delivery systems with advanced 3D models represents a significant step forward in identifying new targets for drug development, evaluating novel therapeutic options, and personalized medicine for endometrial cancer patients.

## 5. Conclusion

This study represents a significant leap in endometrial cancer research, achiving three major milestones: the development of a specialized 3D model of endometrial cancer, the creation of a novel nanoparticle delivery system for Paclitaxel, and the comprehensive preclinical high-throughput drug screening of this innovative chemotherapy treatment using our model. The results of this study underscore the utility of our 3D model for providing a nuanced understanding of endometrial cancer, demonstrating its capability to accurately replicate the complex interplay of cancerous and endothelial cells, and highlighting its potential as a valuable tool for advanced cancer research and therapeutic development.

## Supplementary information

**Figure 1S.**
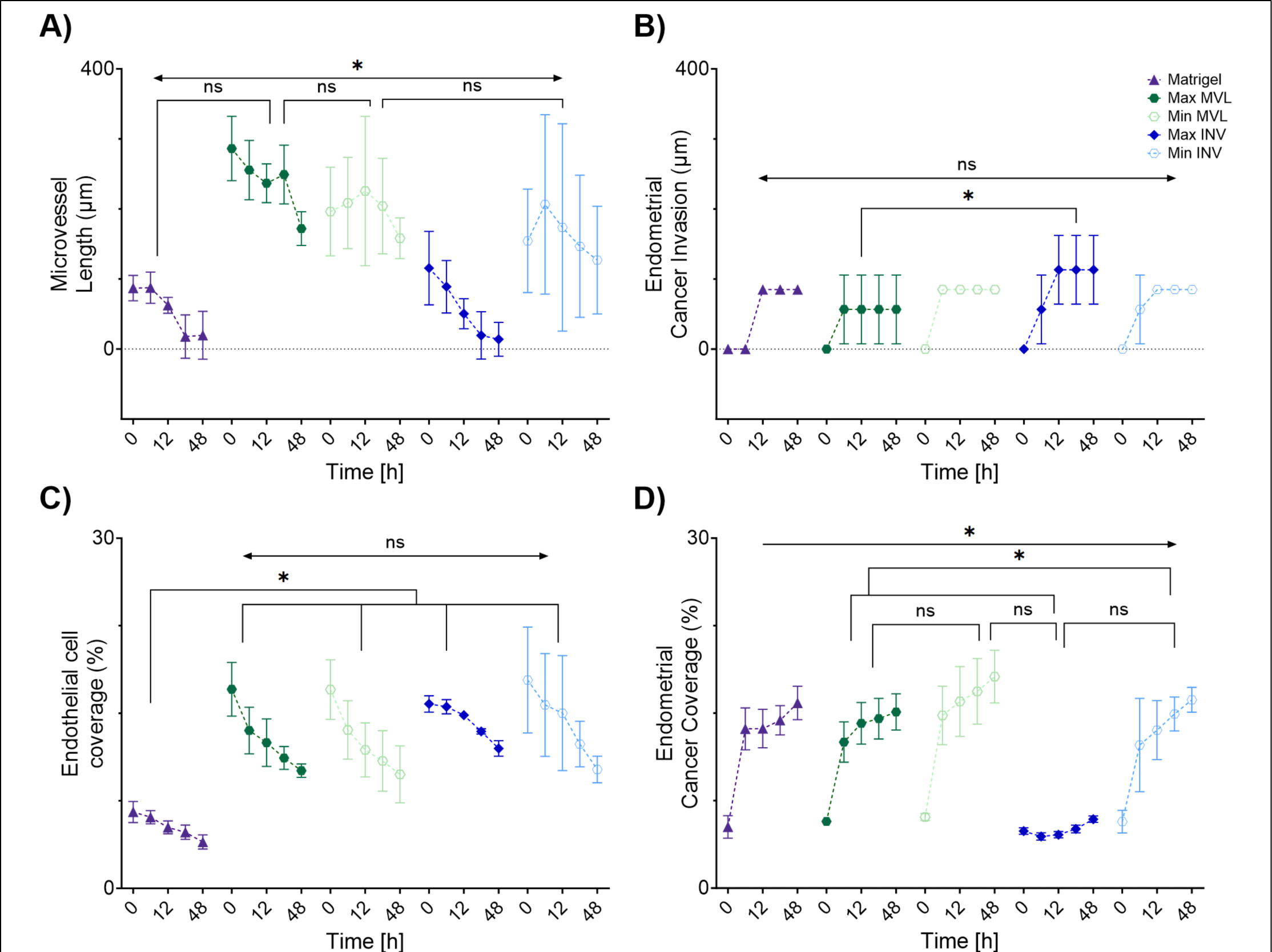
Validation of the DOE model. The desirability function algorithm was used to identify the hydrogel formulations predicted to maximize and minimize microvessel length (MVL) and endometrial cancer invasion (INV) separately. Phenotypic responses, **(A)** microvessel length **(B)** endometrial cancer invasion (**C**) endothelial cell coverage, and (**E**) endometrial cancer coverage were measured every 3 hours for 48 hours. Matrigel was used as a control. * *p* <0.05 compared with the mean of each group. Data were analyzed using a two-way ANOVA with Tukey post-test. Data represent the mean ± SD (n = 4).

**Figure 2S.**
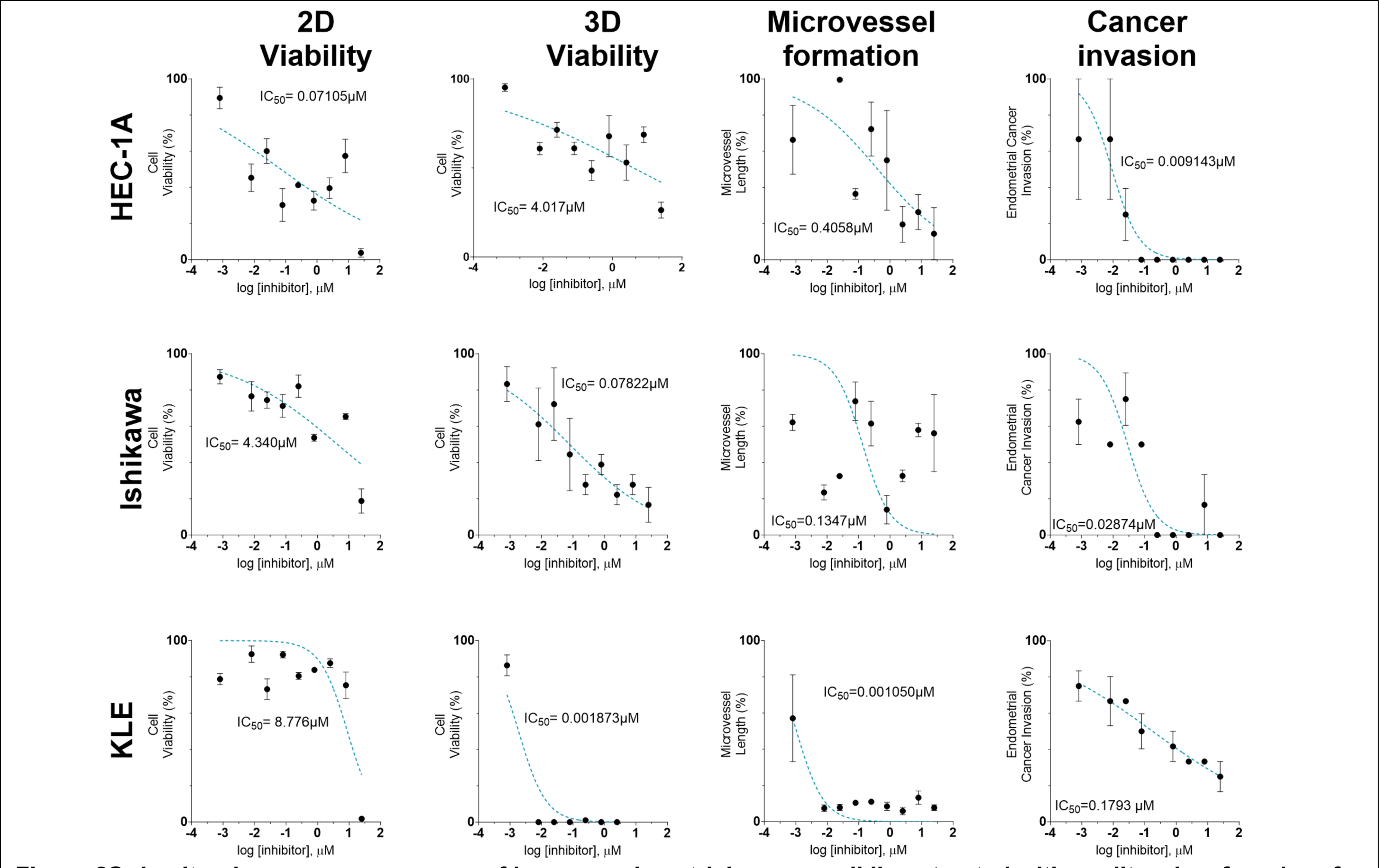
*In vitro* dose response curves of human endometrial cancer cell lines treated with paclitaxel as free drug for 24 hours. Cancer cells were seeded in 96 well plates either on TCP or co-cultured with endothelial cells (hMVEC) in 3D using our optimized hydrogel formulation. Cells were treated with 0.008 – 25 µM of paclitaxel and cell response was evaluated at 24 hours post dosage. Each cell response is normalized to the average of the values observed in the absence of drug. Cell viability and phenotypic cell response were measured in **(A)** HEC-1A **(B)** Ishikawa, and (**C**) KLE cells. Data represent the mean ± SD (n = 4).

**Figure 3S.**
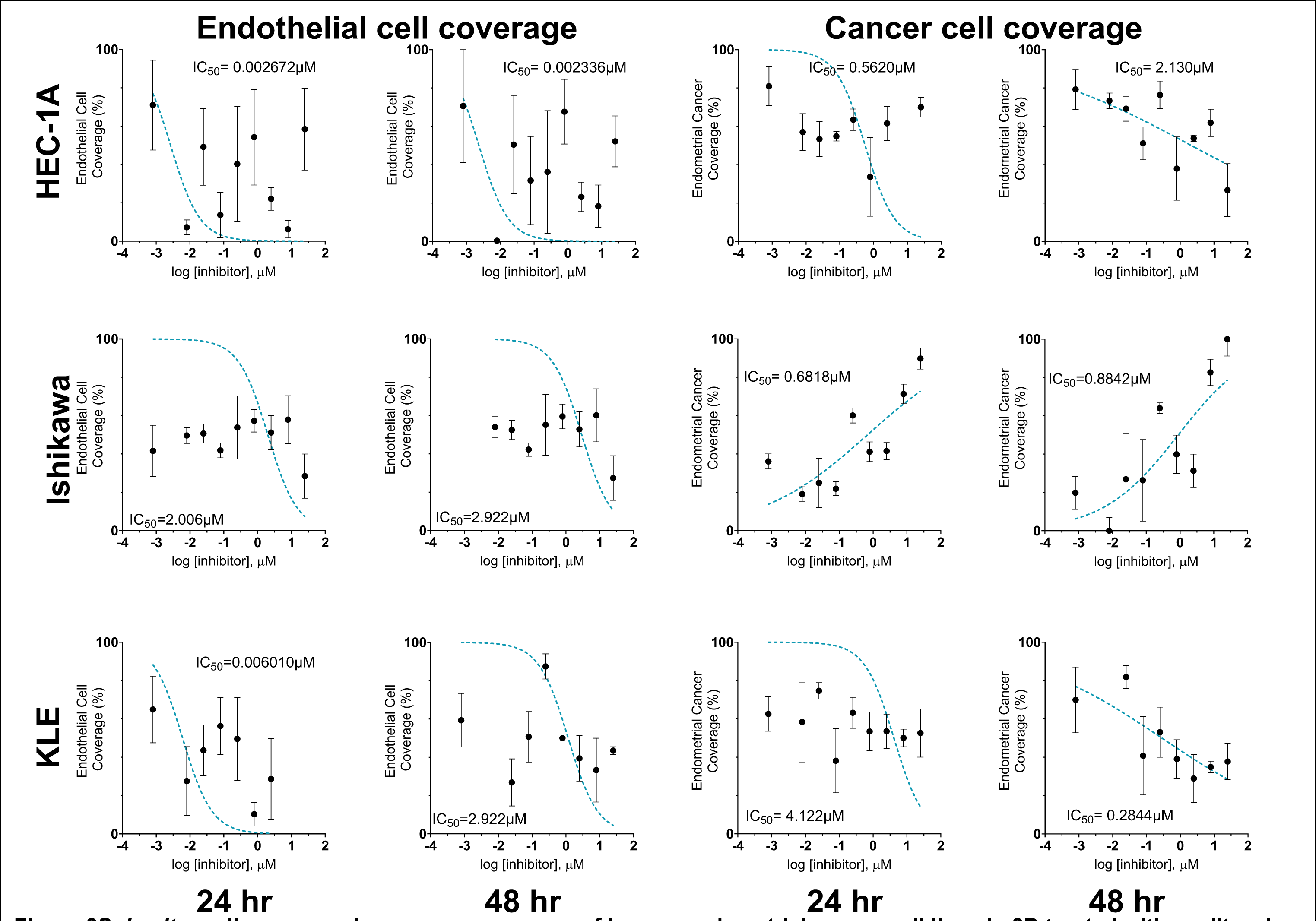
*In vitro* cell coverage dose response curves of human endometrial cancer cell lines in 3D treated with paclitaxel as free drug over time. Cancer cells were seeded in 96 well plates and co-cultured in the 3D multilayer model with hMVEC cells. Plates were treated with 0.008 – 25 µM of paclitaxel and cell response was evaluated at 24 hours and 48 hours post dosage. Each cell response is normalized to the average of the values observed in the absence of drug. Cancer cell coverage and hMVEC coverage were measured in the co-culture models with **(A)** HEC-1A **(B)** Ishikawa, and (**C**) KLE cells. Data represent the mean ± SD (n = 4).

**Figure 4S.**
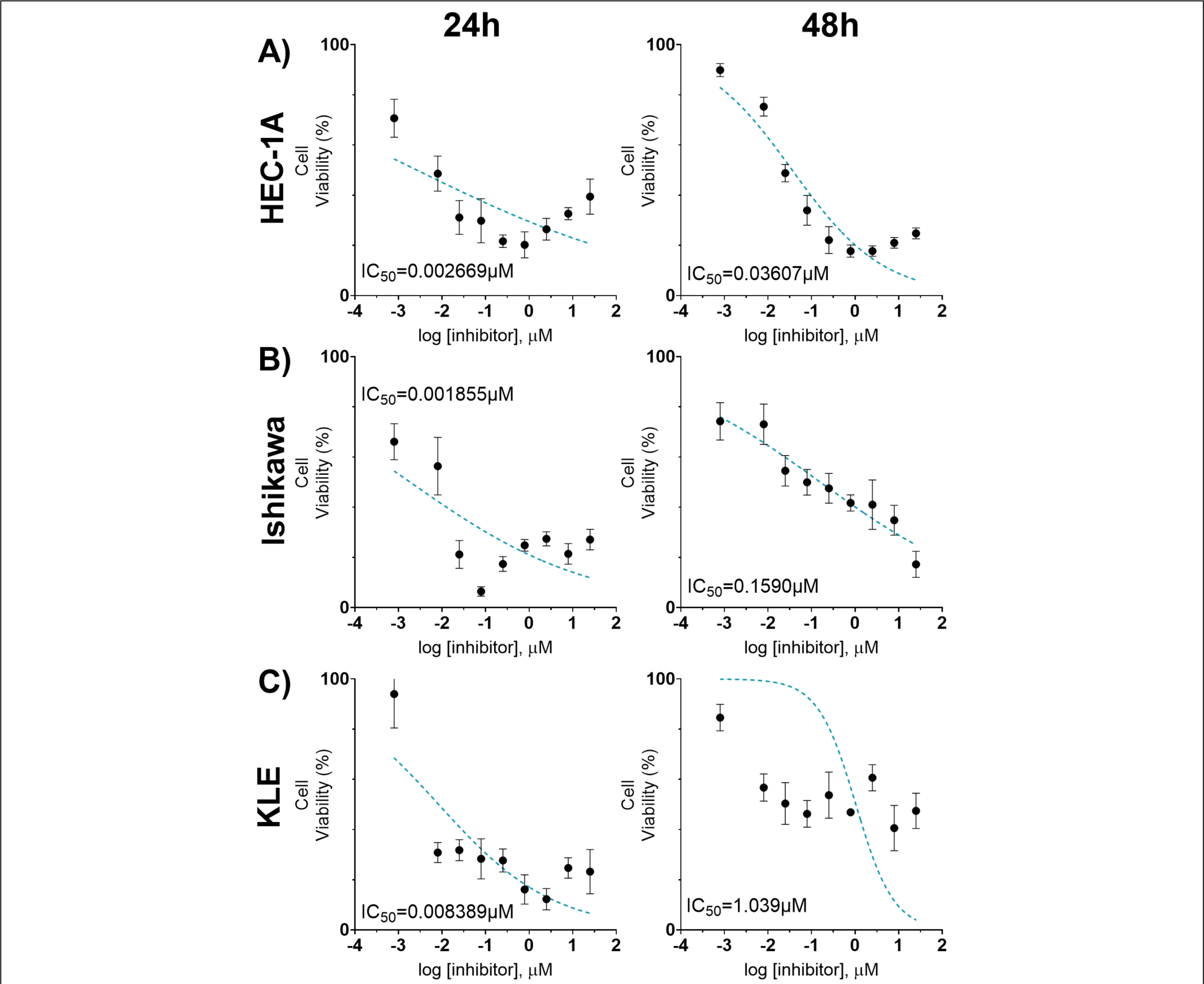
*In vitro* dose response curves of human endometrial cancer cell lines in 2D treated with paclitaxel encapsulated in PCL. Cell viability was compared between endometrial cancer cell lines (HEC-1A, Ishikawa, KLE) seeded in 96 well tissue culture plates (2D) when they were treated with paclitaxel encapsulated in polycaprolactone nanoparticles (PCL). Cells were cultured for 24 hours, then treated with PCL nanoparticles for 48 hours. By measuring absorbance from MTT dye, we recorded the cell viability after 24 hours and 48 hours of exposure to the drug. Cells were treated with 0.008 – 25 µM of drug dispense on top. Each cell response is normalized to the average of the values observed in the absence of drug. Cell viability was quantified in **(A)** HEC-1A, **(B)** Ishikawa, and **(C)** KLE cells. Data represent the mean ± SD (n = 4). IC50 values are presented on each graph.

**Supplemental Table 1.** Design criteria for the Design of the experiment model. Input variables and range are listed.

| Input variable | Abbr. | Units | Type | Range |
| --- | --- | --- | --- | --- |
| Collagen IV | Col<sub>T</sub> | mg/mL | Quantitative | 0.01 to 0.2 |
| Fibronectin | FN<sub>T</sub> | mg/mL | Quantitative | 0.125 to 0.175 |
| Fibrinogen | FG<sub>B</sub> | mg/mL | Quantitative | 0.5 to 2.5 |
| Collagen - Bottom | Col<sub>B</sub> | mg/mL | Quantitative | 0.5 to 2.5 |
| Laminin | Lam | ug/mL | Quantitative | 0.5 to 2 |
| Synthetic Polymer - Top | Syn<sub>T</sub> |  | Qualitative | PEGDA,<br>GelMA |
| Synthetic Polymer - Bottom | Syn<sub>B</sub> |  | Qualitative | PEGDA,<br>GelMA |

<sub>T</sub> and <sub>B</sub> correspond to the top and bottom hydrogel layers.

**Supplemental Table 2.** Hydrogel formulations predicted to maximize or minimize cell phenotypes of interest in the bottom hydrogel (B) or top hydrogel (T). FG = fibrinogen, Syn = synthetic polymer, FN = fibronectin, LMN = laminin.

| Response | [Col I]<sub>B</sub><br>(mg/mL) | [FG]<sub>B</sub><br>(mg/mL) | [Syn]<sub>B</sub> | [Col IV]<sub>T</sub><br>(mg/mL) | [FN]<sub>T</sub><br>(mg/mL) | [LMN]<sub>T</sub><br>(μg/mL) | [Syn]<sub>T</sub> |
| --- | --- | --- | --- | --- | --- | --- | --- |
| Maximize endothelial microvessel length | 1.80 | 1.35 | PEGDA | 0.20 | 0.13 | 2.00 | PEGDA |
| Maximize cancer invasion | 1.80 | 1.92 | GelMA | 0.008 | 0.18 | 2.00 | GelMA |
| Minimize endothelial microvessel length | 1.92 | 0.56 | GelMA | 0.20 | 0.13 | 0.56 | PEGDA |
| Minimize cancer invasion | 2.50 | 0.50 | GelMA | 0.20 | 0.13 | 0.53 | PEGDA |

**Supplemental Table 3.** Input variable combinations examined in D-optimal DOE design.

| Exp No | Exp Name | Run Order | Incl/Excl | Collagen Top | Fibronectin Top | Fibrinogen Bottom | Collagen Bottom | Synthetic Polymer Top | Synthetic Polymer Bottom | Laminin Top |
| --- | --- | --- | --- | --- | --- | --- | --- | --- | --- | --- |
| 1 | N1 | 36 | Incl | 0.2 | 0.125 | 0.5 | 2.5 | PEGDA | PEGDA | 0.5 |
| 2 | N2 | 40 | Incl | 0.01 | 0.125 | 0.5 | 0.5 | PEGDA | PEGDA | 2 |
| 3 | N3 | 30 | Incl | 0.01 | 0.125 | 2.5 | 1.83333 | PEGDA | PEGDA | 0.5 |
| 4 | N4 | 16 | Incl | 0.01 | 0.175 | 0.5 | 2.5 | PEGDA | PEGDA | 1 |
| 5 | N5 | 43 | Incl | 0.01 | 0.175 | 2.5 | 1.16667 | PEGDA | PEGDA | 2 |
| 6 | N6 | 29 | Incl | 0.01 | 0.175 | 1.16667 | 0.5 | PEGDA | PEGDA | 0.5 |
| 7 | N7 | 8 | Incl | 0.2 | 0.175 | 0.5 | 1.16667 | PEGDA | PEGDA | 0.5 |
| 8 | N8 | 35 | Incl | 0.2 | 0.175 | 1.83333 | 0.5 | PEGDA | PEGDA | 2 |
| 9 | N9 | 33 | Incl | 0.2 | 0.15833 | 0.5 | 2.5 | PEGDA | PEGDA | 2 |
| 10 | N10 | 31 | Incl | 0.2 | 0.14167 | 2.5 | 0.5 | PEGDA | PEGDA | 0.5 |
| 11 | N11 | 1 | Incl | 0.136667 | 0.125 | 2.5 | 2.5 | PEGDA | PEGDA | 2 |
| 12 | N12 | 26 | Incl | 0.2 | 0.175 | 2.5 | 2.5 | GELMA | PEGDA | 0.5 |
| 13 | N13 | 11 | Incl | 0.01 | 0.125 | 2.5 | 2.5 | GELMA | PEGDA | 2 |
| 14 | N14 | 17 | Incl | 0.01 | 0.125 | 1.83333 | 0.5 | GELMA | PEGDA | 0.5 |
| 15 | N15 | 39 | Incl | 0.01 | 0.175 | 0.5 | 2.5 | GELMA | PEGDA | 1.5 |
| 16 | N16 | 21 | Incl | 0.01 | 0.175 | 2.5 | 0.5 | GELMA | PEGDA | 1 |
| 17 | N17 | 3 | Incl | 0.01 | 0.14167 | 0.5 | 2.5 | GELMA | PEGDA | 0.5 |
| 18 | N18 | 34 | Incl | 0.2 | 0.125 | 0.5 | 1.83333 | GELMA | PEGDA | 2 |
| 19 | N19 | 9 | Incl | 0.2 | 0.125 | 2.5 | 0.5 | GELMA | PEGDA | 1.5 |
| 20 | N20 | 24 | Incl | 0.136667 | 0.125 | 0.5 | 0.5 | GELMA | PEGDA | 0.5 |
| 21 | N21 | 23 | Incl | 0.073333 | 0.175 | 0.5 | 0.5 | GELMA | PEGDA | 2 |
| 22 | N22 | 38 | Incl | 0.01 | 0.125 | 0.5 | 2.5 | PEGDA | GELMA | 0.5 |
| 23 | N23 | 2 | Incl | 0.2 | 0.175 | 2.5 | 2.5 | PEGDA | GELMA | 0.5 |
| 24 | N24 | 6 | Incl | 0.01 | 0.125 | 2.5 | 2.5 | PEGDA | GELMA | 2 |
| 25 | N25 | 37 | Incl | 0.01 | 0.125 | 2.5 | 0.5 | PEGDA | GELMA | 1 |
| 26 | N26 | 28 | Incl | 0.01 | 0.175 | 0.5 | 1.83333 | PEGDA | GELMA | 2 |
| 27 | N27 | 27 | Incl | 0.01 | 0.15833 | 0.5 | 0.5 | PEGDA | GELMA | 0.5 |
| 28 | N28 | 41 | Incl | 0.2 | 0.125 | 2.5 | 1.16667 | PEGDA | GELMA | 2 |
| 29 | N29 | 5 | Incl | 0.2 | 0.125 | 1.16667 | 0.5 | PEGDA | GELMA | 0.5 |
| 30 | N30 | 25 | Incl | 0.2 | 0.175 | 0.5 | 0.5 | PEGDA | GELMA | 1.5 |
| 31 | N31 | 44 | Incl | 0.073333 | 0.175 | 2.5 | 0.5 | PEGDA | GELMA | 2 |
| 32 | N32 | 32 | Incl | 0.01 | 0.125 | 0.5 | 0.5 | GELMA | GELMA | 0.5 |
| 33 | N33 | 19 | Incl | 0.2 | 0.175 | 2.5 | 0.5 | GELMA | GELMA | 0.5 |
| 34 | N34 | 12 | Incl | 0.2 | 0.175 | 0.5 | 2.5 | GELMA | GELMA | 0.5 |
| 35 | N35 | 7 | Incl | 0.2 | 0.125 | 2.5 | 2.5 | GELMA | GELMA | 0.5 |
| 36 | N36 | 4 | Incl | 0.01 | 0.175 | 2.5 | 2.5 | GELMA | GELMA | 0.5 |
| 37 | N37 | 14 | Incl | 0.2 | 0.175 | 2.5 | 2.5 | GELMA | GELMA | 2 |
| 38 | N38 | 15 | Incl | 0.01 | 0.175 | 1.16667 | 0.5 | GELMA | GELMA | 2 |
| 39 | N39 | 18 | Incl | 0.2 | 0.14167 | 0.5 | 0.5 | GELMA | GELMA | 2 |
| 40 | N40 | 22 | Incl | 0.073333 | 0.125 | 0.5 | 2.5 | GELMA | GELMA | 2 |
| 41 | N41 | 45 | Incl | 0.136667 | 0.125 | 2.5 | 0.5 | GELMA | GELMA | 2 |
| 42 | N42 | 42 | Incl | 0.105 | 0.15 | 1.5 | 1.5 | GELMA | GELMA | 1.25 |
| 43 | N43 | 20 | Incl | 0.105 | 0.15 | 1.5 | 1.5 | GELMA | GELMA | 1.25 |
| 44 | N44 | 10 | Incl | 0.105 | 0.15 | 1.5 | 1.5 | GELMA | GELMA | 1.25 |
| 45 | N45 | 13 | Incl | 0.105 | 0.15 | 1.5 | 1.5 | GELMA | GELMA | 1.25 |

## References

1. Padežnik, T., Oleksy, A., Cokan, A., Takač, I. & Sobočan, M. Changes in the Extracellular Matrix in Endometrial and Cervical Cancer: A Systematic Review. Int J Mol Sci 24, 5463 (2023).

2. Rowlands, C. E. et al. Particles and Prejudice: Nanomedicine Approaches to Reducing Health Disparities in Endometrial Cancer. Small e2300096 (2023) doi:10.1002/smll.202300096.

3. O’Donnell, J. et al. Ipatasertib exhibits anti-tumorigenic effects and enhances sensitivity to paclitaxel in endometrial cancer in vitro and in vivo. Int J Oncol 63, 103 (2023).

4. Chitcholtan, K., Sykes, P. H. & Evans, J. J. The resistance of intracellular mediators to doxorubicin and cisplatin are distinct in 3D and 2D endometrial cancer. J Transl Med 10, 38 (2012).

5. Halla, K. Emerging Treatment Options for Advanced or Recurrent Endometrial Cancer. J Adv Pract Oncol 13, 45–59 (2022).

6. Alqahtani, F. Y., Aleanizy, F. S., El Tahir, E., Alkahtani, H. M. & AlQuadeib, B. T. Chapter Three - Paclitaxel. in Profiles of Drug Substances, Excipients and Related Methodology vol. 44 205–238 (Academic Press, 2019).

7. McMeekin, D. S. et al. The relationship between histology and outcome in advanced and recurrent endometrial cancer patients participating in first-line chemotherapy trials: a Gynecologic Oncology Group study. Gynecol Oncol 106, 16–22 (2007).

8. Ebeid, K. et al. Synthetically lethal nanoparticles for treatment of endometrial cancer. Nat Nanotechnol 13, 72–81 (2018).

9. Manning, A. N., Rowlands, C. E., Saindon, H. & Givens, B. E. Tuning the Emulsion Properties Influences the Size of Poly(Caprolactone) Particles for Drug Delivery Applications. AAPS J 25, 100 (2023).

10. Arruebo, M. et al. Assessment of the evolution of cancer treatment therapies. Cancers (Basel) 3, 3279–3330 (2011).

11. Soppimath, K. S., Aminabhavi, T. M., Kulkarni, A. R. & Rudzinski, W. E. Biodegradable polymeric nanoparticles as drug delivery devices. Journal of Controlled Release 70, 1–20 (2001).

12. Espinoza, S. M., Patil, H. I., San Martin Martinez, E., Casañas Pimentel, R. & Ige, P. P. Poly-ε-caprolactone (PCL), a promising polymer for pharmaceutical and biomedical applications: Focus on nanomedicine in cancer. International Journal of Polymeric Materials and Polymeric Biomaterials 69, 85–126 (2020).

13. Ma, P. & Mumper, R. J. Paclitaxel Nano-Delivery Systems: A Comprehensive Review. J Nanomed Nanotechnol 4, 1000164 (2013).

14. Chang, S. H., Lee, H. J., Park, S., Kim, Y. & Jeong, B. Fast Degradable Polycaprolactone for Drug Delivery. Biomacromolecules 19, 2302–2307 (2018).

15. Chitcholtan, K., Asselin, E., Parent, S., Sykes, P. H. & Evans, J. J. Differences in growth properties of endometrial cancer in three dimensional (3D) culture and 2D cell monolayer. Exp Cell Res 319, 75–87 (2013).

16. Ahn, J. et al. Three-dimensional microengineered vascularised endometrium-on-a-chip. Hum Reprod 36, 2720–2731 (2021).

17. Jamaluddin, M. F. B. et al. Bovine and human endometrium-derived hydrogels support organoid culture from healthy and cancerous tissues. Proc Natl Acad Sci U S A 119, e2208040119 (2022).

18. Huang, Y. et al. Mimicking the Endometrial Cancer Tumor Microenvironment to Reprogram Tumor-Associated Macrophages in Disintegrable Supramolecular Gelatin Hydrogel. Int J Nanomedicine 15, 4625–4637 (2020).

19. Cadena, I. A. et al. Engineering high throughput screening platforms of cervical cancer. Journal of Biomedical Materials Research Part A 111, 747–764 (2023).

20. Benton, G., Arnaoutova, I., George, J., Kleinman, H. K. & Koblinski, J. Matrigel: from discovery and ECM mimicry to assays and models for cancer research. Adv Drug Deliv Rev 79–80, 3–18 (2014).

21. Aisenbrey, E. A. & Murphy, W. L. Synthetic alternatives to Matrigel. Nat Rev Mater 5, 539– 551 (2020).

22. Differential sensitivity to paclitaxel-induced apoptosis and growth suppression in paclitaxel-resistant cell lines established from HEC-1 human endometrial adenocarcinoma cells. https://www.spandidos-publications.com/10.3892/ijo.2012.1600.

23. Liu, S. & Li, X. Autophagy inhibition enhances sensitivity of endometrial carcinoma cells to paclitaxel. Int J Oncol 46, 2399–2408 (2015).

24. Dong, P. et al. Long Non-Coding RNA TMPO-AS1 Promotes GLUT1-Mediated Glycolysis and Paclitaxel Resistance in Endometrial Cancer Cells by Interacting With miR-140 and miR-143. Frontiers in Oncology 12, (2022).

25. Yanokura, M., Banno, K. & Aoki, D. MicroRNA-34b expression enhances chemosensitivity of endometrial cancer cells to paclitaxel. International Journal of Oncology 57, 1145–1156 (2020).

26. Engineering high throughput screening platforms of cervical cancer - Cadena - 2023 - Journal of Biomedical Materials Research Part A - Wiley Online Library. https://onlinelibrary.wiley.com/doi/abs/10.1002/jbm.a.37522.

27. Zhong, L. et al. Small molecules in targeted cancer therapy: advances, challenges, and future perspectives. Signal Transduct Target Ther 6, 201 (2021).

28. Sanoufi, M. R., Aljaberi, A., Hamdan, I. & Al-Zoubi, N. The use of design of experiments to develop hot melt extrudates for extended release of diclofenac sodium. Pharmaceutical Development and Technology 25, 187–196 (2020).

29. McCoy, M. G., Seo, B. R., Choi, S. & Fischbach, C. Collagen I hydrogel microstructure and composition conjointly regulate vascular network formation. Acta Biomater 44, 200–208 (2016).

30. Kroon, M. E., van Schie, M. L. J., van der Vecht, B., van Hinsbergh, V. W. M. & Koolwijk, P. Collagen type 1 retards tube formation by human microvascular endothelial cells in a fibrin matrix. Angiogenesis 5, 257–265 (2002).

31. Tanaka, R. et al. Three-dimensional coculture of endometrial cancer cells and fibroblasts in human placenta derived collagen sponges and expression matrix metalloproteinases in these cells☆. Gynecologic Oncology 90, 297–304 (2003).

32. Ueda, M. et al. [In vitro study on the effect of sex steroid and growth factor on growth and laminin, collagen IV, and tissue plasminogen activator production of normal endometrial cells and endometrial cancer cells in culture]. Nihon Sanka Fujinka Gakkai Zasshi 44, 1219– 1226 (1992).

33. Van Sinderen, M., Griffiths, M., Menkhorst, E., Niven, K. & Dimitriadis, E. Restoration of microRNA-29c in type I endometrioid cancer reduced endometrial cancer cell growth. Oncol Lett 18, 2684–2693 (2019).

34. Berger, A. J., Linsmeier, K. M., Kreeger, P. K. & Masters, K. S. Decoupling the effects of stiffness and fiber density on cellular behaviors via an interpenetrating network of gelatin-methacrylate and collagen. Biomaterials 141, 125–135 (2017).

35. Yoshida, S. et al. Fibronectin mediates activation of stromal fibroblasts by SPARC in endometrial cancer cells. BMC Cancer 21, 156 (2021).

36. Abbas, Y. et al. Tissue stiffness at the human maternal–fetal interface. Hum Reprod 34, 1999–2008 (2019).

37. Kiss, M. Z. et al. Frequency-dependent complex modulus of the uterus: preliminary results. Phys Med Biol 51, 3683–3695 (2006).

38. Matsuzaki, S., Canis, M., Pouly, J.-L. & Darcha, C. Soft matrices inhibit cell proliferation and inactivate the fibrotic phenotype of deep endometriotic stromal cells in vitro†. Human Reproduction 31, 541–553 (2016).

39. Rowlands, C. E. et al. Particles and Prejudice: Nanomedicine Approaches to Reducing Health Disparities in Endometrial Cancer. Small **n/a**, 2300096.

40. Alhaj-Suliman, S. O., Wafa, E. I. & Salem, A. K. Engineering nanosystems to overcome barriers to cancer diagnosis and treatment. Adv Drug Deliv Rev 189, 114482 (2022).

41. Cucinotto, I. et al. Nanoparticle Albumin Bound Paclitaxel in the Treatment of Human Cancer: Nanodelivery Reaches Prime-Time? J Drug Deliv 2013, 905091 (2013).

42. Tian, Z. & Yao, W. Albumin-Bound Paclitaxel: Worthy of Further Study in Sarcomas. Front Oncol 12, 815900 (2022).

43. Chinnathambi, A., Awad Alahmadi, T. & Ali Alharbi, S. Biogenesis of copper nanoparticles (Cu-NPs) using leaf extract of Allium noeanum, antioxidant and in-vitro cytotoxicity. Artificial Cells, Nanomedicine, and Biotechnology 49, 500–510 (2021).

44. Ebeid, K. et al. Synthetically lethal nanoparticles for treatment of endometrial cancer. Nature Nanotech 13, 72–81 (2018).

45. Stewart, A., Denoyer, D., Gao, X. & Toh, Y.-C. The FDA modernisation act 2.0: Bringing non-animal technologies to the regulatory table. Drug Discov Today 28, 103496 (2023).

